# Repurposing approved protein kinase inhibitors as potent anti-leishmanials targeting *Leishmania* MAP kinases

**DOI:** 10.1101/2024.02.16.580710

**Authors:** Anindita Bhattacharjee, Arka Bagchi, Solanki Sarkar, Sriparna Bawali, Arijit Bhattacharya, Arunima Biswas

## Abstract

Leishmaniasis is a vast array of chronic diseases caused by the protozoan parasite of the genus *Leishmania* transmitted by the sandfly belonging to the genus *Phlebotomus* (Old World) and *Lutzomyia* (New World). Among the various factors that contribute to the virulence and pathogenesis of the disease, protein kinases play crucial role in modulating the parasites’ biology in the host system. There are no specific drugs identified to treat the disease specifically. Moreover, resistance against approved anti-leishmanials has made it important to look for a few pre-clinical candidates against visceral leishmaniasis. MAP kinase is one such candidate of protein kinase family that regulates cell proliferation in eukaryotes. In this study, we have identified 23 MAPKs from *Leishmania* genome and have screened 12 FDA approved protein kinase inhibitors against those MAPKs in search of a potential lead for new drug exploration. Among the inhibitors, sorafenib and imatinib have been identified as potential drug candidates based on multiple criteria including binding affinity, ADME scores, absorption and their ability to cross blood-brain barrier. Furthermore, these drugs showed excellent anti-proliferative effects in *Leishmania* promastigotes, resulted in change of cell morphology, flagellar length, promoted cell cycle arrest through ROS generation and reduced intra-macrophage parasitic burden. Collectively, these results imply involvement of MAP kinases in infectivity and survival of the parasite and can pave the avenue for repurposing sorafenib and imatinib as anti-leishmanial agents.

## 1. Introduction

The protozoan parasites of the genus *Leishmania*, which are responsible for a spectrum of neglected tropical diseases known as leishmaniases, are spread by female sandflies of the genus *Phlebotomus* (old world) and *Lutzomyia* (new world), which serve as the vector of the pathogen. Almost 10–12 million individuals are infected by the *Leishmania* parasite, which results in a variety of illnesses known as leishmaniases. 350 million individuals on an average are at a high risk of *Leishmania* transmission affecting 98 countries globally with an annual report of 2 million new cases. A great diversity of different forms of human leishmaniasis are caused by 20 different species of *Leishmania.* Among the different types of leishmaniasis, visceral leishmaniasis is the most fatal form of the disease caused by *Leishmania donovani* that thrives as promastigote in sandfly gut at 22°C at a m pH 7.4 and at 37°C at a pH 5.5 in the mammalian macrophages where it transforms into amastigotes [1]. Differentiation of promastigotes to amastigotes is a controlled process where *Leishmania* proteins undergo post-translational modifications like methylation, acetylation, glycosylation, fucosylation, and phosphorylation [2]. Phosphorylation in amastigotes occurs in proteins whose biological functions are linked to protein degradation, stress response, and signal transduction associated with the differentiation of the parasite. But the exploration of kinase action is yet to achieve comprehensive knowledge on the role of protein kinases in cell cycle regulation in amastigotes. In the past ten years, some scientific progress has been made in the treatment, diagnosis, and management of leishmaniasis, and the cost of various significant medications has decreased. Miltefosine (Miltex), antimonial sodium stibogluconate (Pentostam), and meglumine antimoniate (Glucantime) are the core elements of traditional chemotherapeutic drugs for leishmaniasis. The chemicals have undesirable side effects, and drug resistance is on the rise. Alternative drugs like paromomycin and amphotericin B, the conventional anti-leishmanial drugs, are also restricted because of their high toxicity and expense [3]. However, there are accounts outlining the hazardous effects of pharmaceuticals on patients, and these drugs are not always readily available or effective. Multidrug resistance (MDR) is becoming more prominent in leishmaniasis [4]. Due to the lack of vaccines, recent research is now focused on identifying new targets and developing substitute medications. However, there are currently no effective vaccines against leishmaniasis inspite of the massive efforts to discover novel drugs and vaccines. Therefore, discovering incredibly potent drugs is the main impediment in the management of leishmaniasis.

Identification of potential drug targets is the first step towards the intricate drug discovery process. A desirable drug target in a pathogen is generally a protein that is required for the pathogen’s survival and is absent in or differs from that of the host homolog. Protein kinases are the key regulators of biological processes and they have drawn a lot of interest as possible therapeutic targets for a spectrum of infectious disorders. *Leishmania* cell division is a very complex process and the existence of different promastigote cell types with visible morphological changes occurring during cell cycle progression adds more complications to the already much complicated process. Thus, cell cycle progression in *Leishmania* and its detailed study is a requirement for comprehensive understanding of the survival and infectivity of the parasites in both the hosts. The biochemical and morphological changes that take place during the life cycle of *Leishmania* are perhaps the consequences of programmed changes in the expression of genes in response to the alterations in the external environment of the parasites [5]. Expression of constitutive as well as stage-specific proteins was revealed by genome-wide time course *L. donovani* differentiation which also established the process to be highly programmed and regulated. Post-translational modification is one of the essential events taking place during *Leishmania* differentiation which includes phosphorylation in both promastigote and amastigote forms. Protein phosphorylation plays an important role in the regulation of cell growth, transformation and differentiation in diverse eukaryotes [6]. Leishmanial protein kinases and phosphatases activity might be regulated during the process of parasite differentiation [7]. In other similar kinetoplastid organisms like *Trypanosoma brucei*, not only serine but also threonine and tyrosine kinase activity have been found to be specifically regulated during development of trypanosome parasites [8], [9], [10]. Few phosphotyrosines are also identified in *Leishmania*, particularly in the proteins that take part in the signal transduction pathway. Furthermore, a total of 28 differentially regulated proteins were identified from a proteome analysis of *Leishmania infantum* promastigote stage for the first time. Apart from this, a comprehensive MS-based phosphorylation mapping of phosphoproteins from *Trypanosoma cruzi* epimastigotes has identified 237 phosphopeptides out of 119 distinct proteins [11]. Moreover, presence of at least 7 tyrosine-phosphorylated proteins was confirmed in *T. cruzi.* The fact that *Leishmania* is phylogenetically close to prokaryotes and contains unique p-sites and motifs that differ from those of higher eukaryotes is intriguing, justifying further in-depth analysis of their phosphoproteome [11]. In another study, 29 protein kinases were identified which are required for promastigote to amastigote differentiation and survival of the parasite in mammalian macrophages. Apart from 13 unknown protein kinases, 5 protein kinases were involved in MAPK signaling, 2 CDKs and 2 casein kinase II were also identified [12]. In *Leishmania*, MAPKs are required for the development of flagella and survival inside the hostile environment of the mammalian macrophage [13], [14], [15], [16], [17]. Among the 179 protein kinases that have been identified in *Leishmania*, 15 genes were established as typical MAPKs [18]. In *L. mexicana*, MAPK1 (LmxMPK1) plays a major role in amastigote survival inside macrophage [19]. LmxMPK3 and LmxMPK9 are associated with flagellar length regulation in promastigotes, whereas LmxMPK4 is involved with transformation and virulence of the parasite [15], [17]. LdMPK1 is associated with the phosphorylation of heat shock proteins (HSPs). Reduction in HSP phosphorylation due to decrease in MAPK1 levels makes them less stable and subsequently subjects them to proteolysis [20]. Moreover, inactivation of LmxMPK1 annihilated parasite virulence during mammalian infection [21]. This altogether suggests that in *Leishmania,* like the higher eukaryotes, MAP kinase pathway is an intrinsic component of the signal transduction and plays crucial role in various cellular functions and infectivity of the parasite.A large data bank is now available for all the protein kinases that compose the *Leishmania* kinome providing an opportunity for deciphering various signaling pathways involved with various important aspects of the parasite’s biology such as stress response, differentiation and replication. Moreover, till January 1, 2024, there are 80 small molecules, which function as inhibitors of mammalian protein kinases, have been approved by Food and Drug Administration (FDA) as therapeutic agents against several infectious, as well as chronic diseases [22]. Therefore, among the many protein kinases present in *L. donovani* kinome, we have studied the interaction of MAP kinases with FDA approved drugs using computational methods which predict protein structures and ligand-protein interactions. MAPKs, being a significant component of the signaling pathway regulating the cell cycle progression and transformation of the parasite from promastigote to amastigote, stand out as one of the potent drug targets for anti-leishmanial compounds. In order to find new inhibitors of *Leishmania* MAPKs, structure-based virtual screening was implemented to anticipate the binding affinity of the FDA-approved drugs. This approach involved computational docking of ligands (inhibitors) with a receptor (LdMAPks), followed by scoring and ranking of ligands to discover potential leads. The complexes predicted by the computational docking studies were further assessed for their stability and dynamics by Molecular Dynamics (MD) simulations, to predict their behavior in cellular conditions. From the molecular docking studies, 2 inhibitors namely, sorafenib and imatinib are selected based on their binding affinity scores and their efficacy is assessed against *Leishmania* promastigotes and amastigotes. A conventional MAPK inhibitor, p38 MAPK inhibitor IV is considered as a positive control. A striking role of MAPK in regulating parasitic morphology is speculated through microscopic studies. Moreover, the inhibitors resulted in decreased cell viability in both promastigote and axenic amastigotes and enhanced ROS generation which validate them as prospective repurposed drugs against *Leishmania*.

## 2. Materials and methods

### 2.1. Ethics Statement

The entire study was in strict accordance with the recommendations in the Guide for the Care and Use of Laboratory Animals of the Committee for the Purpose of Control and Supervision of the Experiments on Animals (CPCSEA). The protocol was approved by the Institutional Animal Ethics Committee on Animal Experiments of the University of Kalyani (892/GO/Re/S/01/CPCSEA).

The maintenance and culture of *L. donovani* in the cell culture facility of the Department of Zoology, University of Kalyani was also in strict regulations according to the recommendations in the Institutional Biosafety Committee (Ref No. IBSC-KU/2023/A4), Kalyani University (IBSC-KU).

### 2.2. Parasites and cell culture

The pathogenic *Leishmania donovani* strain AG83 (MHOM/IN/1993/Ag83) was maintained within BALB/c by administering promastigotes every 6 weeks via tail vein. The promastigotes were obtained after dissecting the mice in sterile condition and then allowing the splenic amastigotes to transform into promastigotes in medium M199 (Thermo Fisher Scientific, Waltham, MA, USA), supplemented with 10% heat-inactivated fetal calf serum (FCS) along with penicillin (50U/ml) and streptomycin (50mg/ml) at 22°C.

Simutaneously, RAW 264.7, a murine macrophage cell line with adherent properties, was grown in RPMI 1640 medium (Invitrogen). The medium was supplemented with 10% fetal calf serum (FCS), 100 U/ml penicillin, and 100 µg/ml streptomycin, and the cells were maintained at 37°C with 5% CO_2_. For *in vivo* infection, female BALB/c mice weighing approximately 20 g were intravenously injected with 1 × 10^7^ stationary phase promastigotes of *L. donovani* through the tail vein. Parasites were retrieved by aseptically extracting the spleen and liver from the infected mice, and the quantification of parasite burdens was determined using Leishman–Donovan units (LDUs). LDUs were calculated by assessing the number of parasites per 1,000 nucleated cells multiplied by the organ weight (g).

### 2.3. Parasite viability assay

To assess parasite viability, the treated and control cells were subjected to a viability assay by incubating them in 0.5 mg/ml 3-(4,5-dimethylthiazol-2-yl)-2,5-diphenyltetrazolium bromide (MTT) for 3 hours. Subsequently, 100 µl of 0.04 N HCl in isopropyl alcohol was added. The MTT assay is based on the principle that living mitochondria convert MTT to a dark blue compound called formazan, which is soluble in acid isopropyl alcohol. The detection of formazan was carried out at 570 nm using a microplate reader. The percentage viability was determined by calculating the ratio of optical density (OD) values in wells with treated cells to the wells with controls, multiplied by 100.

### 2.4. Flow cytometry and cell cycle analysis

To analyse the impact of various inhibitors on the progression of the cell cycle, the quantification of DNA content within the cells was performed. Propidium iodide, a DNA-binding dye, was utilized for this purpose. A 5 ml sample of parasites (0.5–1 × 10^7 cells/ml) was centrifuged at 1,000 g for 10 minutes at 22°C, underwent two washes in PBS, and was then re-suspended in 70% ice-cold methanol for fixation. The fixed cells were stored at −20°C for future use. Subsequently, the cells were treated with 20 mg/ml of RNase A and incubated at 37°C for 1 hour before proceeding to analysis. Propidium iodide was then added for DNA staining, and the DNA content of 20,000 cells was analysed [23]. Flow cytometry was conducted using the BD LSRFortessaTM (BD Biosciences), and the obtained data were analyzed using BD FACSDiva 9.0TM (BD Biosciences).

### 2.5. Macrophage infectivity assay

RAW 264.7 cells were cultured in RPMI-1640 supplemented with 10% FCS. After counting, the cells were centrifuged at 1,500 rpm for 10 minutes at 4°C, resuspended in culture medium, and distributed over 18 mm^2^ coverslips, followed by overnight incubation. Subsequently, the cells were treated with various inhibitors (details provided later) and infected with *L. donovani* promastigotes at a ratio of ten parasites per macrophage. The infection was allowed to proceed overnight.

For quantifying the parasite number within the macrophages, the cells were fixed in methanol and stained with DAPI (4′,6-diamidino-2-phenylindole) at a concentration of 1 µg/ml in PBS containing 10 µg/ml of RNase A (Sigma-Aldrich). Visualization of cells was performed using an Olympus IX 81 microscope with an FV1000 confocal system, utilizing a 100× oil immersion Plan Apo (N.A. 1.45) objective. Analysis of the images was conducted using Olympus Fluoview Software (version 3.1a, Tokyo, Japan).

### 2.6. H_2_DCFDA staining and Intracellular ROS measurements

H_2_DCFDA is a cell-permeable dye that becomes fluorescent upon reacting with reactive oxygen species (ROS). In this experiment, *L. donovani* promastigotes were cultured, and log-phase promastigotes were treated with inhibitory doses (IC50) of the specified inhibitors for 24 hours. Following the removal of the media, the cells were washed with PBS and treated with 1 mM H_2_DCFDA dye for 30 minutes. The cells were then incubated at 37°C in the dark. After incubation, the cells were washed twice with PBS and examined under a microscope (20x, Zeiss Axioscope A1) to detect fluorescence-emitting cells. The presence of an intense fluorescence signal indicates the formation of reactive oxygen species (ROS). This method serves to visualize and assess the extent of ROS formation in the treated promastigotes.

### 2.7. Protein structure modelling and analysis

23 Ld MAPK protein structures were built using Swiss-Modelers for all the LdMAPKs after the sequences have been retrieved from Tritrypdb database. The superfamily domain was taken for searching template in Swiss-Modelers. The MAPKs protein structures were selected based on sequence coverage with the template and GMQE scores for the protein models generated by the Swiss-Model and analysed by Ramachandran Plot server where models with 0-1 amino acids were selected for further studies.

### 2.8. Molecular docking studies

Before the docking studies, the ligands (12 FDA-approved inhibitors and 1 conventional MAPK inhibitor) were prepared in OpenBabel software for conversion into PDB format, after the 3D structures of all the ligands were obtained from PubChem [24]. The input file format of proteins and ligand for the molecular docking studies and molecular dynamic simulation is Protein Data Bank (PDB) format. The unnecessary atoms, such as solvent atoms, were removed from these PDB by PyMOL (Schrӧdinger). The molecular docking studies of all the proteins with the inhibitors was performed using the virtual screening tool PyRx, which utilizes autodock vina to identify compounds with good binding scores. Separate structure files for proteins for both the protein (macromolecule) and ligand (micromolecules) were provided to the software in PDB format. The software then prepares the inputs by loading force fields. Then the software was allowed to run independent docking in every site predicted by the Autodock Vina. The output was recorded in form of PDB of the poses of the protein-ligand complexes along with their binding affinity in kcal/mol, estimated Ki and ligand efficiency.

### 2.9. Molecular dynamics simulation

Molecular Dynamics (MD) simulations were carried out for observing the dynamic behaviour of the proteins and protein-ligand complexes and how they evolve with time. GROMACS (version 2022.4) software was utilized in a terascale high-performance computing cluster installed at the S.N. Bose Innovation Centre of the University of Kalyani. Molecular coordinates of the best docked complexes were used for simulating in GROMACS software for 100 nanoseconds (ns). The output trajectories were utilized to compute the root mean square deviation (RMSD), radius of gyration (Rg), root mean square fluctuations (RMSF). The number of stable hydrogen bonds formed between the protein and the ligand throughout the course of simulation were also computed using tool internal to GROMACS.

### 2.10. Evaluation and comparison of drug-likeness and toxicity prediction

The structures of all 12 FDA-approved protein kinase inhibitors were obtained from PubChem in canonical SMILES (Simplified Molecular Input Line Entry System) format, which is a chemical notation consisting of linear text format describing connectivity and chirality of a molecule. These structures were subsequently used to investigate their absorption, distribution, metabolism, excretion, and toxicity (ADMET) properties using the SwissADME platform. (Website link) and ADMETlab 2.0 (Website link). SwissADME is a freely available tool that assesses various physicochemical properties, including the n-octanol/water partition coefficient (log Po/w). This coefficient is a crucial parameter in drug discovery, design, and development, as highlighted in the study by Diana et al. Additionally, SwissADME offers a boiled egg plot, aiding in predicting the substance’s potential for absorption in the gastrointestinal tract and its ability to penetrate the blood-brain barrier. The ADMETlab 2.0 web-based tool is a multi-task graph attention framework equipped with an array of high-quality prediction models

The drug-likeness of these compounds was assessed using Lipinski’s Rule of Five, which outlines key criteria for a compound to be considered likely for use as a drug. According to Lipinski’s rule, a compound is favorable for drug development if it meets the following criteria: no more than 5 hydrogen bond donors, no more than 10 hydrogen bond acceptors, a molecular weight not exceeding 500 KDa, and an octanol/water partition coefficient not exceeding 5. These criteria, in conjunction with other pharmacokinetic parameters, contribute to evaluating the drug-like properties of the compounds.

### 2.11. Statistical Analysis

All the results shown are representatives of a number of independent experiments as mentioned in the figure legends. Statistical differences among the sets of data were determined by Student’s t-test with a P-value <0.05 considered to be significant.

## 3. Results

### 3.1. Identification of 23 MAPK from *Leishmania* genome and virtual screening of FDA approved kinase inhibitors against *Leishmania* MAPKs: Modelling and structure validation

Since MAPKs has a crosstalk with cAMP signaling as obtained from previous data and they play vital role in cell cycle progression and proliferation in the parasite [25], we sought to understand the role of *Leishmania* MAPKs as major protein kinases associated with the cell cycle regulation, infectivity and thus, disease manifestation. 23 MAPKs were identified from *Leishmania* genome according to annotation in TritrypDB database (Table 1) and sequences were retrieved for homology modelling. Homology models of the catalytic domains of LdMAPKs were built using SWISS-MODEL and MODELLER. The protein structures were ranked on the basis GMQE scores and DOPE score for SWISS-MODEL and MODELLER respectively. The models were validated through Ramachandran plot as generated by PROCHECK (in supplementary material). Models with <1% amino acids in the disallowed region were selected for subsequent studies (Figure S1).

**Table 1:**
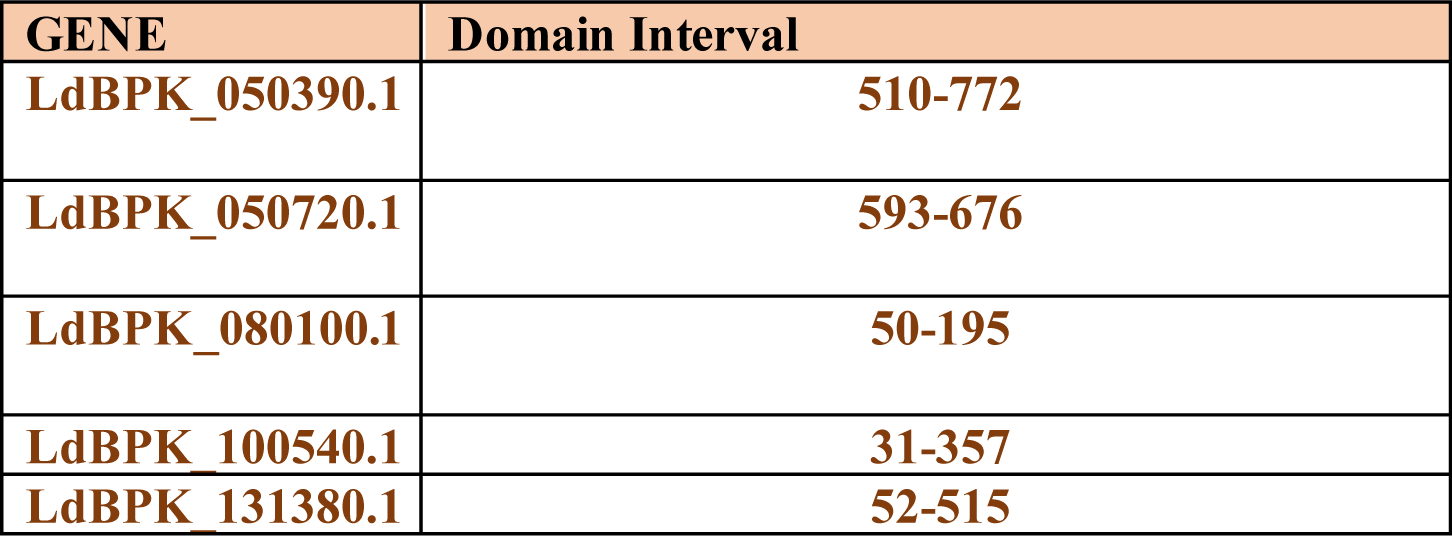
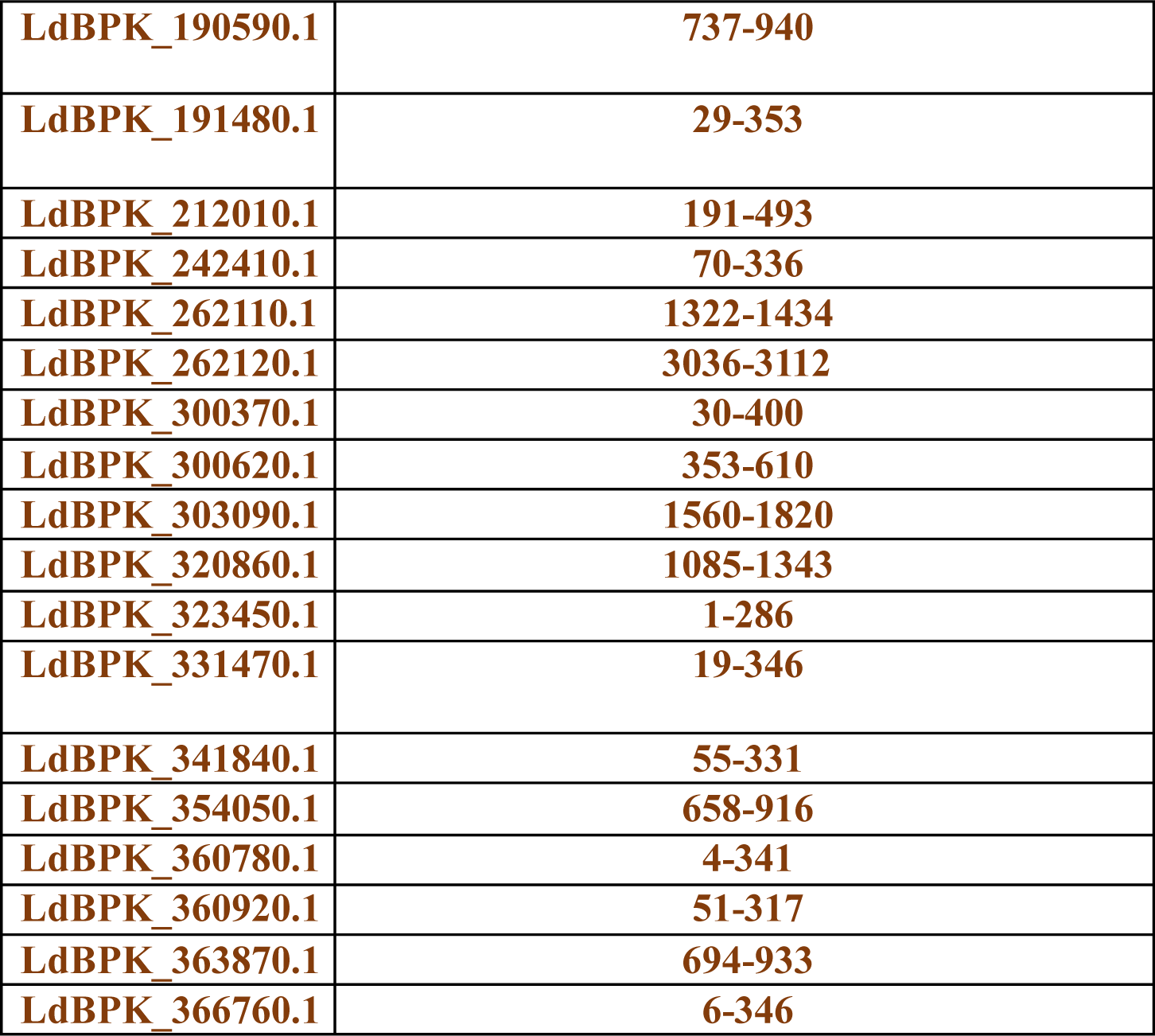
23 MAPKs were identified from genome-wide identification of kinases of MAPK family in *L. donovani*.

A number of MAPK inhibitors have been approved for the treatment of a variety of malignancies and others directed against inflammatory diseases [26], [27]. To identify candidate repurposable kinase inhibitors for *Leishmania* MAPKs, structure-based virtual screening was performed to predict the binding affinity of 12 such FDA-approved inhibitors (enlisted in Table 2) for the MAPK orthologues. The ligand (drugs) structures were retrieved from PubChem.

**Table 2:**
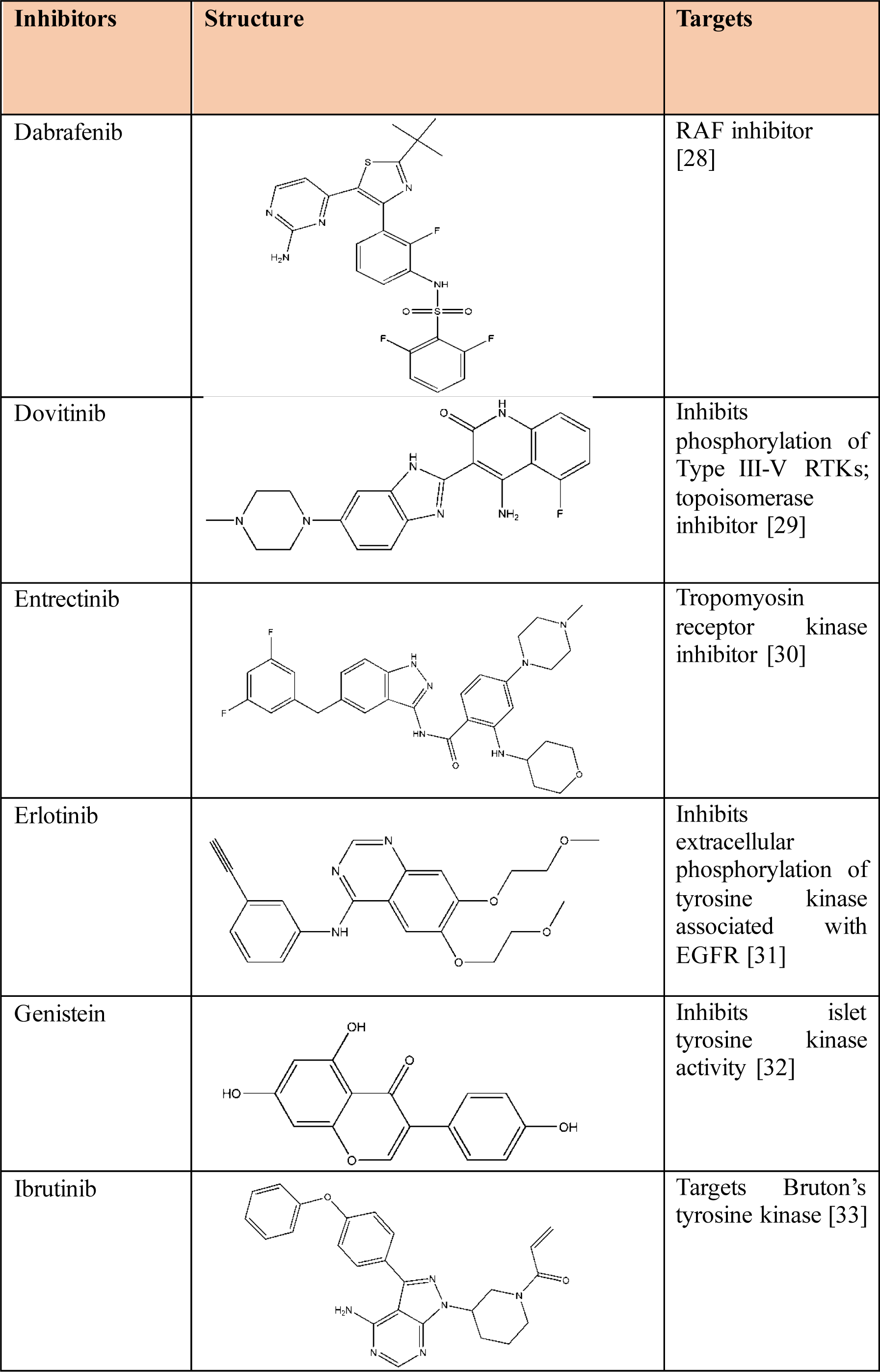
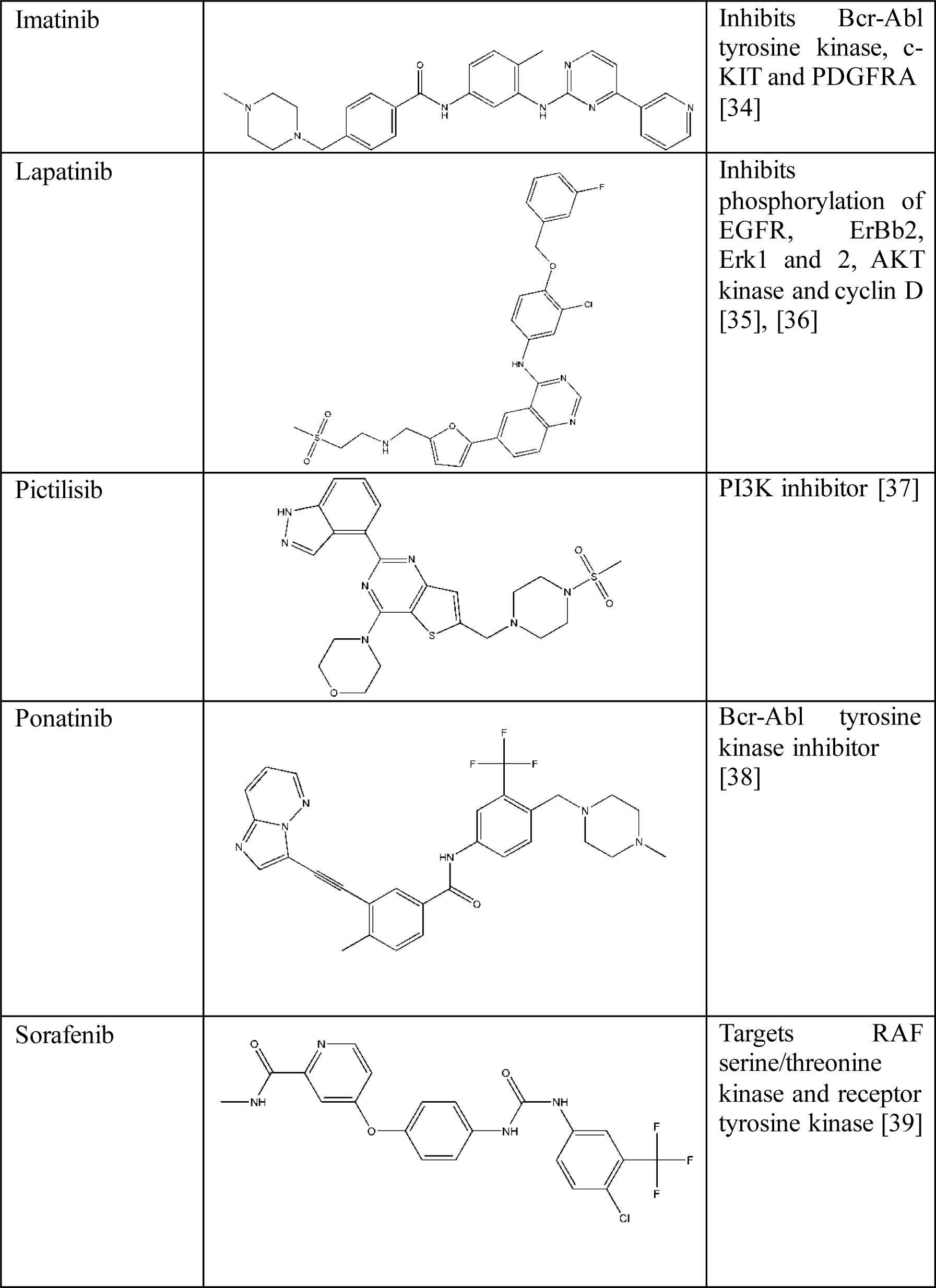
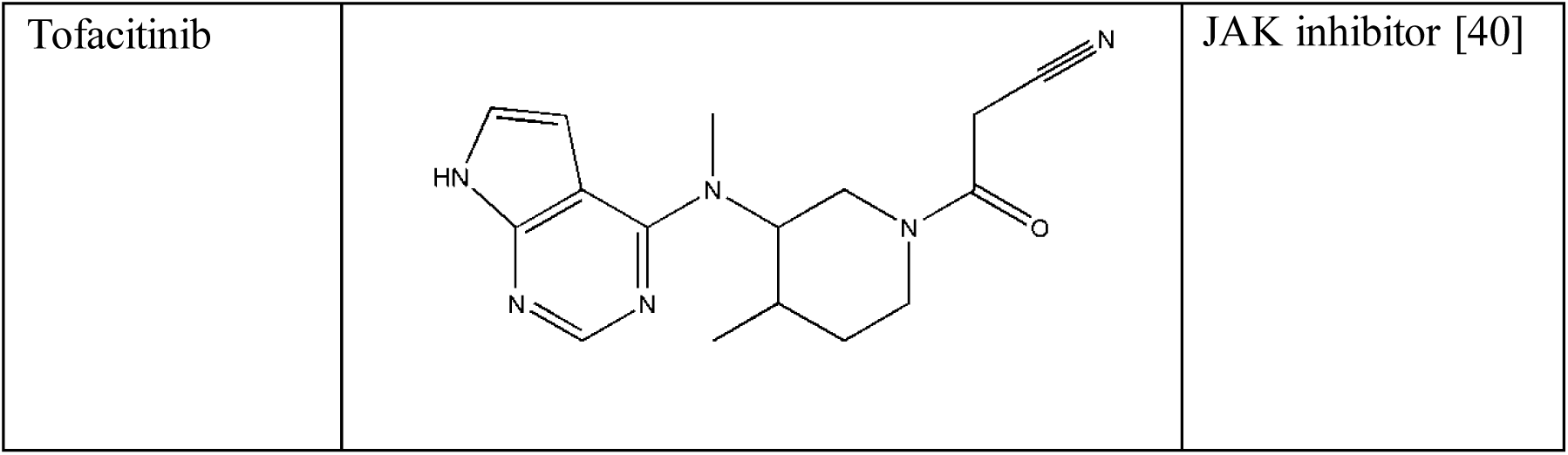
List of FDA-approved protein kinase inhibitors to be used for molecular docking with *Leishmania* MAPKs, their structure and respective targets.

| Inhibitors | Structure | Targets |
| --- | --- | --- |
| Dabrafenib |  | RAF inhibitor [28] |
| Dovitinib |  | Inhibits phosphorylation of Type III-V RTKs; topoisomerase inhibitor [29] |
| Entrectinib |  | Tropomyosin receptor kinase inhibitor [30] |
| Erlotinib |  | Inhibits extracellular phosphorylation of tyrosine kinase associated with EGFR [31] |
| Genistein |  | Inhibits islet tyrosine kinase activity [32] |
| Ibrutinib |  | Targets Bruton's tyrosine kinase [33] |
| Imatinib |  | Inhibits Bcr-Abl tyrosine kinase, c-KIT and PDGFRA [34] |
| Lapatinib |  | Inhibits phosphorylation of EGFR, ErbB2, Erk1 and 2, AKT kinase and cyclin D [35], [36] |
| Pictilisib |  | PI3K inhibitor [37] |
| Ponatinib |  | Bcr-Abl tyrosine kinase inhibitor [38] |
| Sorafenib |  | Targets RAF serine/threonine kinase and receptor tyrosine kinase [39] |
| Tofacitinib |  | JAK inhibitor [40] |

### 3.2. Drug docking and binding analysis

Rigid docking performed using Autodock Vina provided an insight into the binding affinity and hydrogen bond formation of the ligands on the protein kinases. The virtual screening tool PyRx was used to identify compounds withy good binding scores. The structures of the prepped proteins were docked with the ligands into druggable sites to identify best fits. The results of the drug screening are summarized in Table 3 and Figure 1. From the binding energy estimated from 23 dockings for each inhibitor, ligands demonstrating high affinity to the *Leishmania* MAPK orthologues were prioritized based on a hierarchical clustering analysis.

**Figure 1:**
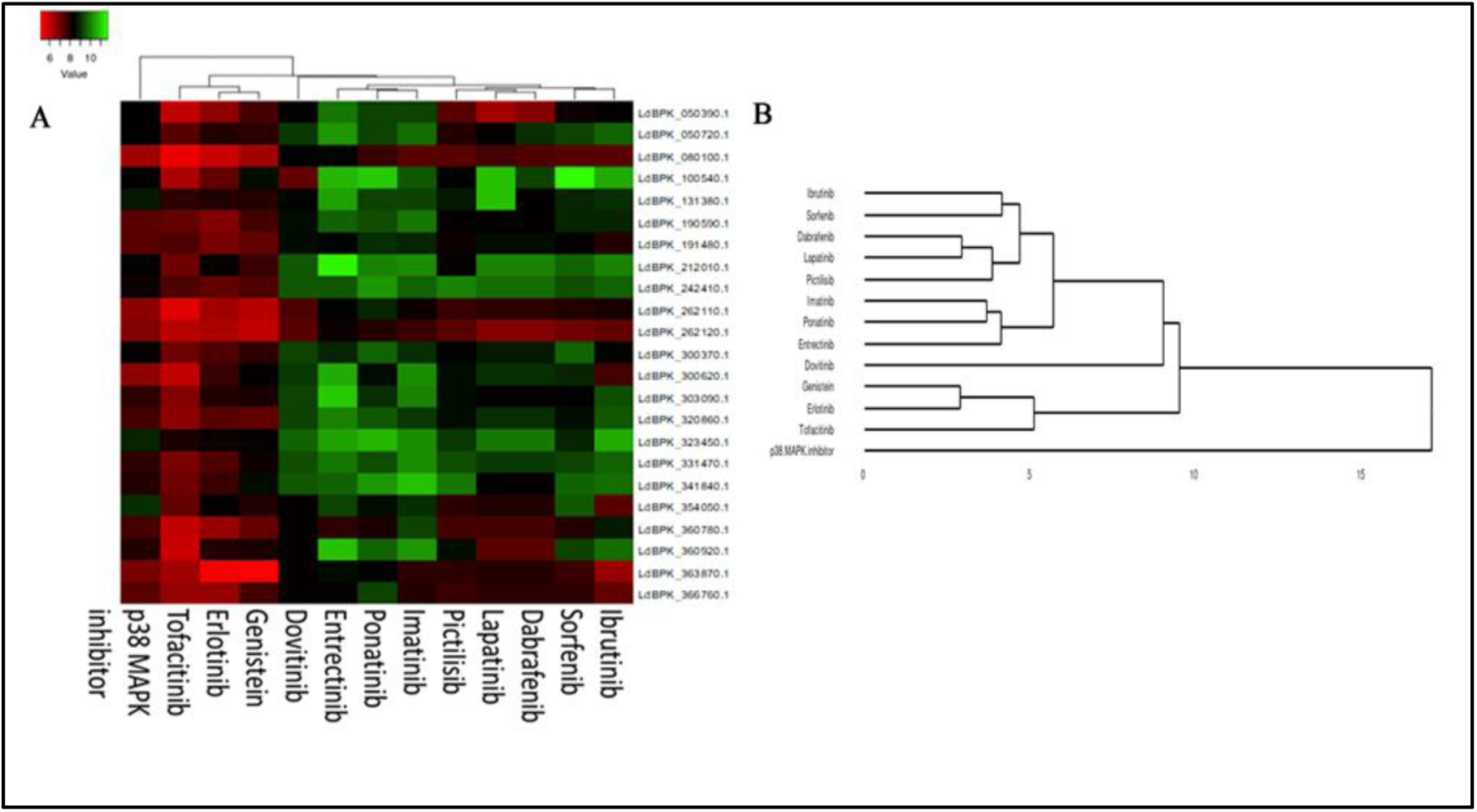
Clustering heatmap of the docking scores of the 13 inhibitors with target proteins is shown. The intensity of color indicates docking score value, green: lowest, red: highest (A). Average linkage clustering obtained from the heatmap. The graphical presentation is representative of high, medium and low efficacy cluster (B).

**Table 3:** Binding energies of all the 23 *Leishmania* MAPKs were docked with the FDA-approved kinase inhibitors along with p38 MAP kinase inhibitor as a control. (Visual representations of the two best fit docked complexes are provided in supplementary material (Figure S3).

| *Leishmania* MAPK kinase genes | p38 MAPK IV inhibitor | Ponatinib | Pictilisib | Lapatinib | Dabrafenib | Sorfenib |
| --- | --- | --- | --- | --- | --- | --- |
| LdBPK_050390.1_Leishmania donovani BPK282A1 | -8.4 | -9.3 | -7.3 | -6.4 | -6.7 | -8.2 |
| LdBPK_050720.1_Leishmania donovani BPK282A1 | -7.2 | -9.3 | -8 | -8.4 | -9 | -9.2 |
| LdBPK_080100.1_Leishmania donovani BPK282A1 | -6.4 | -7.7 | -7.3 | -7.6 | -7.4 | -7.3 |
| LdBPK_100540.1_Leishmania donovani BPK282A1 | -6.9 | -11 | -8.5 | -10.8 | -9.3 | -11.7 |
| LdBPK_131380.1_Leishmania donovani BPK282A1 | -7.7 | -9.3 | -8.7 | -10.8 | -8.4 | -8.9 |
| LdBPK_190590.1_Leishmania donovani BPK282A1 | -7.3 | -9.4 | -8.3 | -8.4 | -8.4 | -8.9 |
| LdBPK_191480.1_Leishmania donovani BPK282A1 | -7.3 | -9 | -8.2 | -8.6 | -8.6 | -8.3 |
| LdBPK_212010.1_Leishmania donovani BPK282A1 | -9.2 | -10 | -8.4 | -10 | -10 | -9.7 |
| LdBPK_242410.1_Leishmania donovani BPK282A1 | -8.2 | -10.3 | -10 | -9.8 | -9.8 | -9.4 |
| LdBPK_262110.1_Leishmania donovani BPK282A1 | -6.6 | -8.8 | -7.7 | -7.8 | -7.8 | -7.9 |
| LdBPK_262120.1_Leishmania donovani BPK282A1 | -6.8 | -7.9 | -7.3 | -6.8 | -6.8 | -7 |
| LdBPK_300370.1_Leishmania donovani BPK282A1 | -8.3 | -9.7 | -8.5 | -8.7 | -8.7 | -9.6 |
| LdBPK_300620.1_Leishmania donovani BPK282A1 | -6.6 | -8.6 | -8.6 | -9 | -9 | -8.9 |
| LdBPK_303090.1_Leishmania donovani BPK282A1 | -7.8 | -9.7 | -8.6 | -8.3 | -8.3 | -8.5 |
| LdBPK_320860.1_Leishmania donovani BPK282A1 | -7.5 | -9 | -8.6 | -9 | -9.2 | -8.6 |
| LdBPK_323450.1_Leishmania donovani BPK282A1 | -8.9 | -10.7 | -9.1 | -9.9 | -9.7 | -8.9 |
| LdBPK_331470.1_Leishmania donovani BPK282A1 | -7.8 | -9.4 | -9.4 | -9.1 | -9 | -9.3 |
| LdBPK_341840.1_Leishmania donovani BPK282A1 | -8 | -10.5 | -9.9 | -8.3 | -8.5 | -9.7 |
| LdBPK_354050.1_Leishmania donovani BPK282A1 | -9 | -8.6 | -7.8 | -8 | -8.1 | -9.5 |
| LdBPK_360780.1_Leishmania donovani BPK282A1 | -7.5 | -8.1 | -7.6 | -7.5 | -7.5 | -7.9 |
| LdBPK_360920.1_Leishmania donovani BPK282A1 | -7.9 | -9.6 | -8.6 | -7.3 | -7.6 | -9.3 |
| LdBPK_363870.1_Leishmania donovani BPK282A1 | -6.9 | -8.5 | -7.8 | -7.9 | -7.4 | -7.7 |
| LdBPK_366760.1_Leishmania donovani BPK282A1 | -7.3 | -9.2 | -7.7 | -7.8 | -7.8 | -7.8 |

| *Leishmania* MAPK kinase genes | Imatinib | Dovitinib | Erlotinib | Entrectinib | Genistein | Tofacitinib | Ibrutinib |
| --- | --- | --- | --- | --- | --- | --- | --- |
| LdBPK_050390.1_Leishmania donovani BPK282A1 | -9.3 | -8.3 | -6.6 | -9.9 | -7.5 | -5.9 | -8.4 |
| LdBPK_050720.1_Leishmania donovani BPK282A1 | -9.8 | -9.1 | -7.9 | -10.3 | -7.8 | -7.3 | -9.7 |
| LdBPK_080100.1_Leishmania donovani BPK282A1 | -7.3 | -8.7 | -5.8 | -8.5 | -6.4 | -5.4 | -7.3 |
| LdBPK_100540.1_Leishmania donovani BPK282A1 | -9.5 | -7.2 | -7.2 | -10.7 | -8.6 | -6.2 | -10.6 |
| LdBPK_131380.1_Leishmania donovani BPK282A1 | -9.3 | -8.5 | -7.9 | -10.5 | -7.9 | -7.8 | -9 |
| LdBPK_190590.1_Leishmania donovani BPK282A1 | -9.9 | -8.6 | -6.8 | -9.7 | -7.5 | -7.2 | -8.9 |
| LdBPK_191480.1_Leishmania donovani BPK282A1 | -8.9 | -8.6 | -6.9 | -8.5 | -7.2 | -7.4 | -7.9 |
| LdBPK_212010.1_Leishmania donovani BPK282A1 | -10.2 | -9.5 | -8.4 | -11.5 | -7.7 | -7 | -10 |
| LdBPK_242410.1_Leishmania donovani BPK282A1 | -9.7 | -9.5 | -7.2 | -9.5 | -7.4 | -7.4 | -9.7 |
| LdBPK_262110.1_Leishmania donovani BPK282A1 | -8.2 | -7.4 | -6.4 | -8.5 | -6.1 | -5.6 | -8.1 |
| LdBPK_262120.1_Leishmania donovani BPK282A1 | -7.7 | -7.3 | -6.2 | -8.2 | -6 | -6.1 | -7.1 |
| LdBPK_300370.1_Leishmania donovani BPK282A1 | -9 | -9.3 | -7.4 | -8.9 | -7.8 | -7 | -8.5 |
| LdBPK_300620.1_Leishmania donovani BPK282A1 | -10.2 | -9.1 | -7.7 | -10.6 | -8.3 | -5.9 | -7.6 |
| LdBPK_303090.1_Leishmania donovani BPK282A1 | -10 | -9.2 | -8 | -11 | -8.1 | -6.7 | -9.4 |
| LdBPK_320860.1_Leishmania donovani BPK282A1 | -9.1 | -9.2 | -7.3 | -10.1 | -7.1 | -6.6 | -9.5 |
| LdBPK_323450.1_Leishmania donovani BPK282A1 | -10.2 | -9.7 | -8.2 | -10.5 | -8.3 | -8.1 | -10.6 |
| LdBPK_331470.1_Leishmania donovani BPK282A1 | -10.2 | -9.4 | -7.4 | -9.9 | -8.2 | -6.9 | -9.7 |
| LdBPK_341840.1_Leishmania donovani BPK282A1 | -10.8 | -9.5 | -7.7 | -9.7 | -8.6 | -7 | -9.8 |
| LdBPK_354050.1_Leishmania donovani BPK282A1 | -9 | -8.4 | -8.4 | -9.2 | -8 | -7.1 | -7.3 |
| LdBPK_360780.1_Leishmania donovani BPK282A1 | -9.3 | -8.3 | -6.5 | -7.8 | -7.1 | -6 | -8.7 |
| LdBPK_360920.1_Leishmania donovani BPK282A1 | -10.3 | -8.4 | -8 | -10.9 | -8.1 | -5.8 | -9.8 |
| LdBPK_363870.1_Leishmania donovani BPK282A1 | -7.8 | -8.3 | -5.3 | -8.6 | -5.1 | -6.3 | -6.5 |
| LdBPK_366760.1_Leishmania donovani BPK282A1 | -7.9 | -8.4 | -6.5 | -8.3 | -7.6 | -6.5 | -7.2 |

Sorafenib displayed binding energy as low as -11.7 kcal/mol, followed by entrectinib and imatinib, -11.5 kcal/mol and -10.8 kcal/mol, respectively with LdBPK_323450.1_ *Leishmania donovani* BPK282A1. Considering the three inhibitors as potential leads, which must be tested *in vitro* and *in vivo* experiments to confirm their real potential as anti-leishmanial drugs. Binding energies obtained from virtual screening of Ld MAPK, LdBPK_323450.1 against all the kinase inhibitors are represented in Table 3. Analysis of PK-inhibitor complexes atomistic molecular dynamics (MD) simulations represent a convenient approach for structure-based drug rationalization.

The comprehensive molecular docking analysis prioritized two kinase inhibitors, sorafenib and imatinib, along with p38MAPKI, which were subjected to MD simulation of 100 ns with the kinase active-site demonstrating highest binding affinity. All the analysis displayed stability of the molecule at the active sites albeit the dynamics varied among the complexes (Fig. 3D). For, LdBPK_323450.1_ *Leishmania donovani* BPK282A1, the apo form simulation demonstrated indicated RMSD of 0.15 from the beginning and remained at that level throughout the simulation period (Fig. 3D). The RMSF profile suggested fluctuation of the atoms within 0.2 nm, indicating stability of the molecule and thereby of the catalytic core (Fig. 2D).

**Figure 2:**
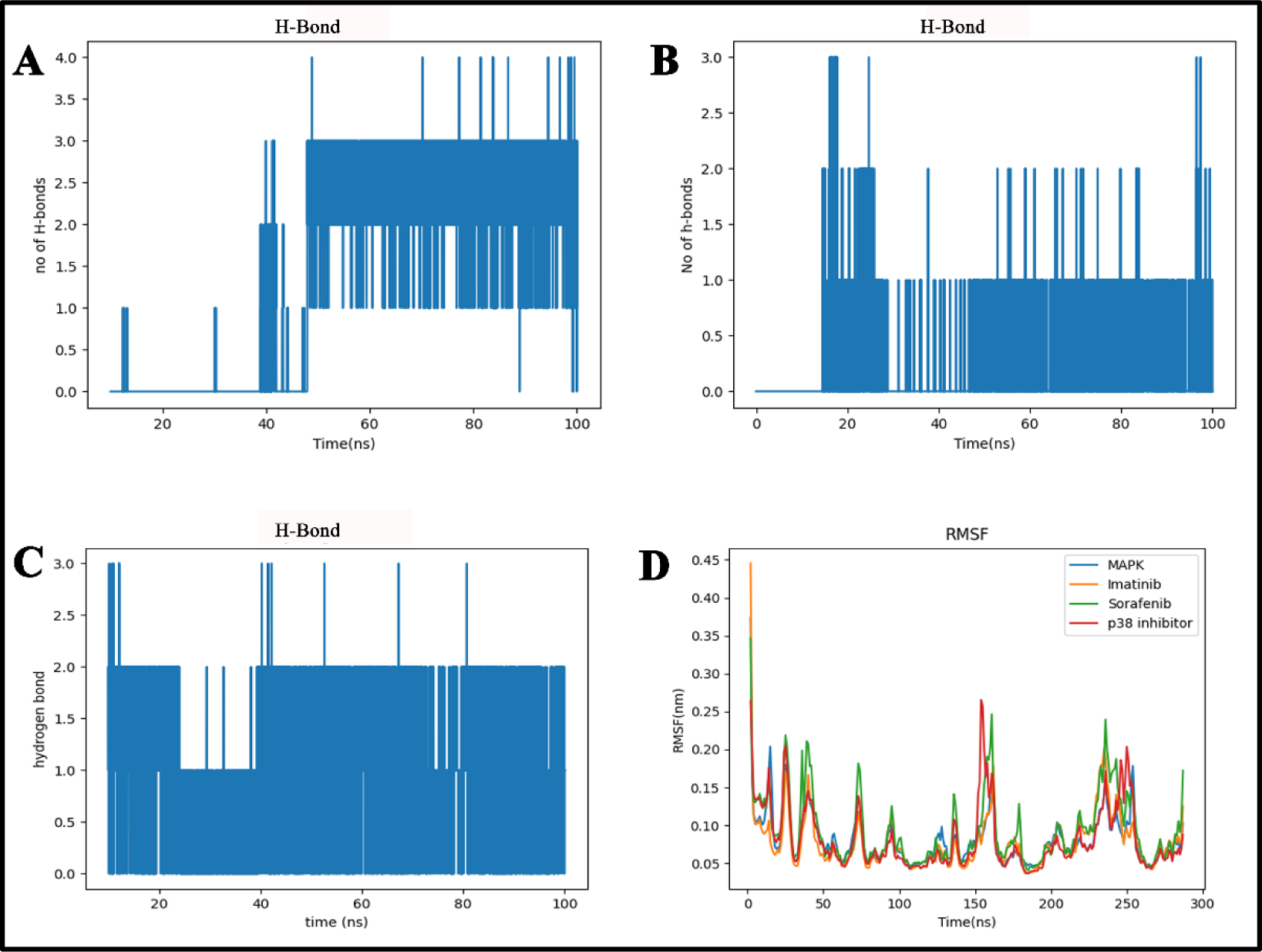
Hydrogen bond formation between the LdBPK_323450.1_ *Leishmania donovani* BPK282A1 and p38 MAPK Inhibitor (A), Sorafenib (B) and Imatinib (C) during the course of simulation is represented. RMSF analysis of 300 amino acid residues is represented for MAPK (blue), p38 MAPK Inhibitor (red), Sorafenib (green) and Imatinib (orange) protein complexes (D).

**Figure 3:**
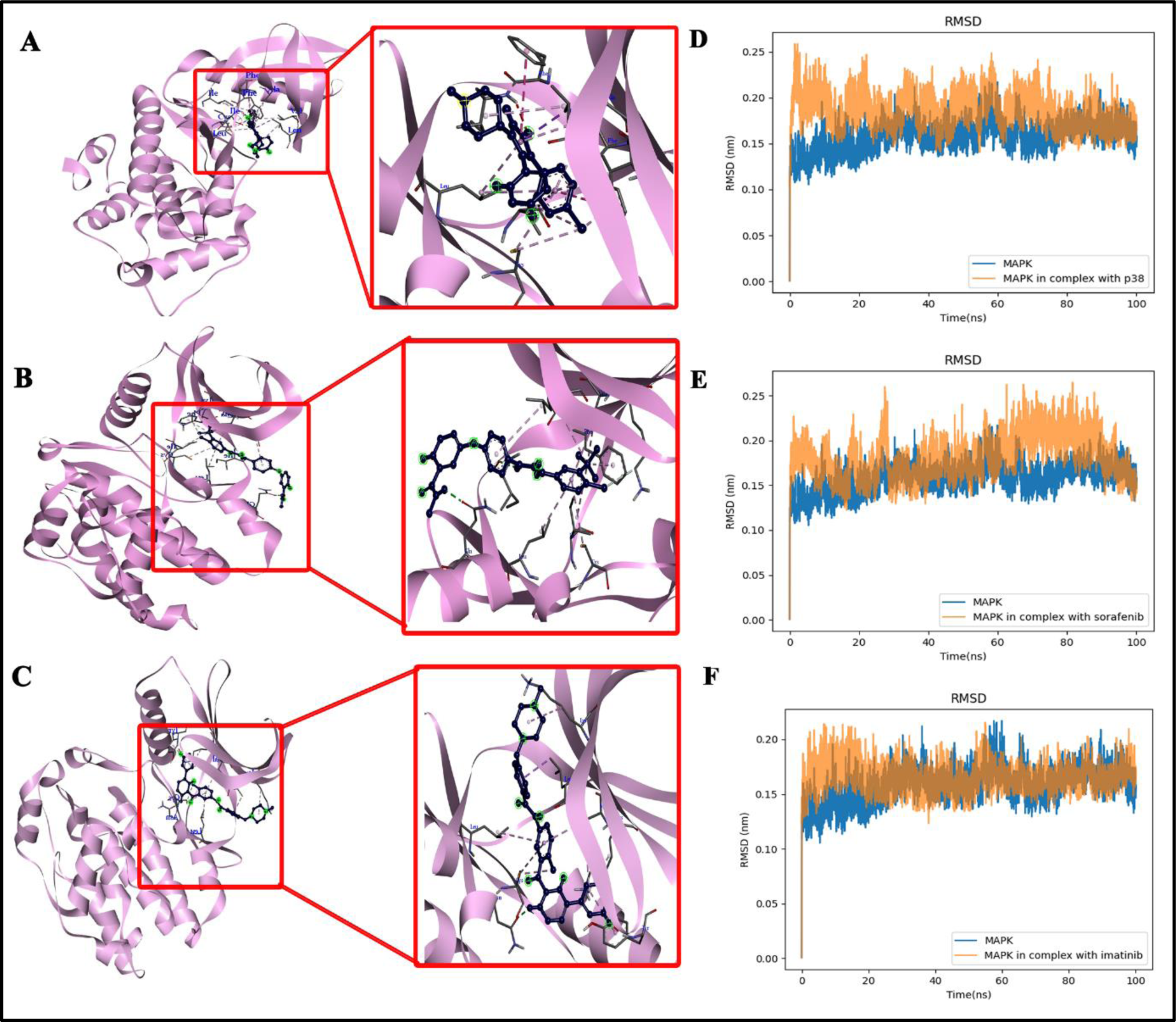
Molecular docking of LdBPK_323450.1_ *Leishmania donovani* BPK282A1 with p38 MAPK Inhibitor (A), Sorafenib (B) and Imatinib (C). (D), (E) and (F) represent the RMSD of MAPK in complex with p38 MAPK Inhibitor, Sorafenib and Imatinib, respectively and comparison with RMSD of MAPK in water for 100 ns MD simulation.

The RMSD of LdBPK_323450.1_ *Leishmania donovani* BPK282A1 with p38 MAPKI projected RMSD between 0.2-0.25 nm for the initial 20 ns, which subsequently converged and stabilized around 0.15 nm link the apo form (Fig. 3D). The RMSF plot suggested fluctuation of residue 150-160 reaching above 0.25 nm, and for 240-250 attaining around 0.2 nm (Fig. 2D). However, the fluctuations were recorded distant from the ligand binding site and the atoms for the residues at the ligand binding site remained stable with restrained side-chain fluctuation throughout. 2-3 stable H-bonds formed between Arg33 and N_4_/N_3_ of p38 MAPKI around 50 ns of the simulation remained stable till the end of the simulation, indicating stable interaction of the protein with p38 MAPKI at the predicted docking site.

As depicted in Fig. 3F, the RMSD of LdBPK_323450.1_ *Leishmania donovani* BPK282A1 in complex with imatinib attained value of 0.2 nm at the initial 17 ns of simulation, however it was stabilized around 0.15 ns like the apo protein subsequently (Fig. 3F). The RMSF plot for the complex indicated a profile similar to the apo form with trivial fluctuations of the atoms of the residues between 220-235 (Fig. 2D). The simulation also indicated formation of 2 H-bonds between Gly88 and imatinib around 40 ns which remained stable till 100 ns (Fig. 2C). The data indicated that imatinib occupied the catalytic domain with least perturbation of the protein’s native conformation.

Simulation for LdBPK_323450.1_ *Leishmania donovani* BPK282A1 in complex with sorafenib displayed RMSD around 0.2 nm till 60 ns with a rise between 60ns and 80 ns to 0.25 nm, and subsequent convergence to 0.15 nm around 90 ns (Fig. 3E). The RMSF profile indicated fluctuation of ∼0.05 to 0.1 nm with respect to apo form for fluctuation of the atoms between residues 30-50, 155-175, and between 225-245 (Fig. 2D). However, the fluctuations did not affect the site for binding with sorafenib. At least one stable H-bond was detected between sorafenib and the kinase which appeared around 47 ns in Leu87 residue and remained stable till 100 ns (Fig. 2B). Collectively, for both imatinib and sorafenib, considerably stable interactions with LdBPK_323450.1_ *Leishmania donovani* BPK282A1 were predicted by the simulation.

### 3.3. Drug-likeness and ADME studies for ligands in comparison to approved anti-leishmanials

Because of the unfavourable pharmacokinetic characteristics and lack of druglikeness, most compounds are unsatisfactory. To ascertain how effectively a chemical is absorbed, distributed, metabolized and excreted out of the body, a pharmacokinetics analysis was carried out. A chemical compound must fulfil certain requirements in order to be considered a potential pharmaceutical. Lipinski’s rule of five is applied to examine a set of characteristics, including molecular weight, lipophilicity, hydrogen bond donors and acceptors. The molecule may be orally accessible if its molecular weight is below 500 Da, there are fewer than 5 hydrogen bonds, fewer than 10 hydrogen acceptors, and the octanol-water partition coefficient LogP is below 5 [41]. The pharmacokinetic and drug-likeness criteria were violated by the majority of the docked ligands, resulting in their rejection as drugs. Among the 12 FDA-approved protein kinase inhibitors, sorafenib and imatinib satisfied Lipinski’s rule of five (Fig. 4E, 4G and 4I and Table 5) along with the conventional p38 MAPK inhibitor. However, Amphotericin B, inspite of being a well-known anti-leishmanial drug, doesn’t fulfil the Lipinski’s rule (Table 5) of five and has low gastrointestinal absorption which makes the discovery of new drugs a necessity.

**Figure 4:**
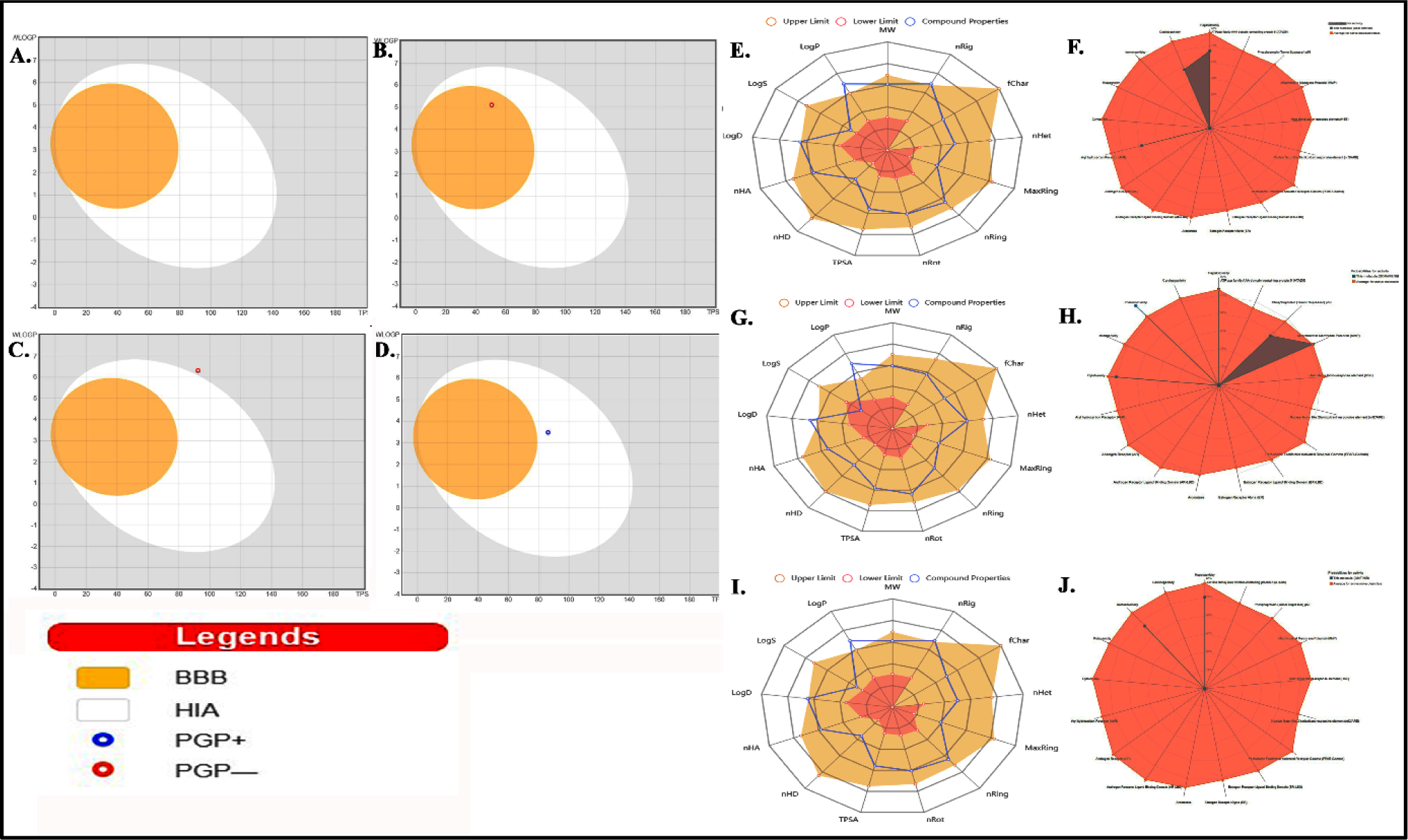
The BOILED-Egg allows for intuitive evaluation of passive gastro-intenstinal absorption (HIA) and brain penetration (BBB) in function of the position of the molecules in the WLOGP-vs-TPSA referential. Positions of p38 MAPKI, sorafenib and imatinib are represented in blot (B-D). Blot (A) shows that the conventional drugs against visceral leishmaniasis, Amphotericin B and Miltefosine, are out of range. Druggability analysis for kinase inhibitors p38 MAPKI, sorafenib and imatinib was analyzed for satisfying Lipinski filter (E, G and I); Toxicity radar for predicting systemic toxicity of the same are also presented separately (F, H and J).

**Table 4:** ADME and drug likeness properties using of Sorafenib and Imatinib along with control drugs, p38 MAPKI, Amphotericin B and Miltefosine using SwissADME.

| Ligand | Molecular weight (g/mol) | Lipophilicity (Log P<sub>o/w</sub> (XLOGP3)) | Hydrogen bond acceptor/donor | Water Solubility (Log S (ESOL)) |
| --- | --- | --- | --- | --- |
| Sorafenib | 464.82 | 4.07 | 7,3 | Moderately soluble |
| Imatinib | 493.60 | 3.52 | 6,2 | Moderately soluble |
| p38 MAPK Inhibitor | 365.79 | 4.40 | 3,1 | Moderately soluble |
| Amphotericin B | 924.08 | 0 | 18,12 | Moderately soluble |
| Miltefosine | 407.57 | 6.79 | 4,0 | Moderately soluble |

**Table 5:** Tabular representation of pharmacokinetic properties of sorafenib and imatinib along with the control drugs like p38 MAPKI, Amphotericin B and Miltefosine.

| Ligand | Druglikeness violations (Lipinski, Ghose, Veber, Egan, Muegge) | Bioavailability | Blood Brain Barrier | Gastrointestinal Absorption | Skin permeability (cm/s) |
| --- | --- | --- | --- | --- | --- |
| Sorafenib | Ghose ( 1 violation, WLOGP3>5.6); Egan (1 violation, WLOGP3>5.88)) | 0.55 | No | Low | -6.25 |
| Imatinib | Ghose ( 2 violations, MW>480, MR>130) | 0.55 | No | High | -6.81 |
| p38 MAPK Inhibitor | No violations | 0.55 | Yes | High | -5.41 |
| Amphotericin B | Lipinski ( 3 violations, MW>500: NorO>10, NHorOH>5)<br>Ghose (3 violations, MW>480, MR>130, #atoms>70)<br>Veber ( 1 violation, TPSA>140)<br>Egan (1 violation, TPSA>131.6)<br>Muegge ( 4 violations, MW>600, TPSA>150, H-acc>10, H-don>5) | 0.17 | No | Low | -11.94 |
| Miltefosine | Ghose ( 2 violations, WLOGP>5.6, #atoms>70)<br>Veber (1 violation, Rotors>10)<br>Egan (1 violation, WLOGP>5.88)<br>Muegge ( 2 violations, XLOGP3>5, Rotors>15) | 0.55 | No | Low | -3.97 |

### 3.4. Effects of the kinase inhibitors on cell morphology and cell cycle progression

The selected drugs (sorafenib and imatinib), along with p38 MAPK IV inhibitor as a positive control were evaluated for anti-proliferative activities in *Leishmania* promastigotes by MTT assay. The results showed that each inhibitor had anti-leishmanial properties that were dose-dependent and significantly cytotoxic. A bar graph representing the IC_50_ values for all the compounds against log phase promastigotes is shown in the Fig. 5A. Sorafenib and imatinib were found to be strongly inhibitory with IC_50_ value as low as 3.5 μM and 7.3 μM for promastigotes and 3.22 μM and 6.95 μM for amastigotes, respectively. A conventional MAPK inhibitor, p38 MAPK inhibitor IV, was found to be lesser cytotoxic as compared to the other compounds with IC_50_ value of 33.3 μM for promastigotes and 29.28 μM for amastigotes (Fig. 5A). The EC_50_ values were also determined after exposing the cells to the inhibitors for 24 hours. Both the kinase inhibitors affected growth with the EC_50_ of 0.95 and 1.6 μM in amastigotes for sorafenib and imatinib, respectively (Fig. 5B). Propidium iodide (PI) FACS analysis of DNA synthesis in G_1_/S boundary synchronised *Leishmania* promastigotes after release from hydroxyurea and treated with the kinase inhibitors was used to investigate whether the cytotoxic effects of the compounds are promoting cell cycle arrest. As shown in the Figure 5C, treatment of *Leishmania* promastigotes with all the inhibitors at their IC_50_ doses resulted in increase in percentage of Sub G_0_/G_1_ cell population and a concomitant decrease in G_0_/G_1_ cell population, indicating a probable apoptosis-like event. The result shows that 3.9% of cells were in Sub G_0_/G_1_ phase in untreated cells, but the percentage increased to 4.8%, 6% and 5.7% in p38 MAPK inhibitor IV, sorafenib and imatinib treated cells, respectively (Fig. 5C).

**Figure 5:**
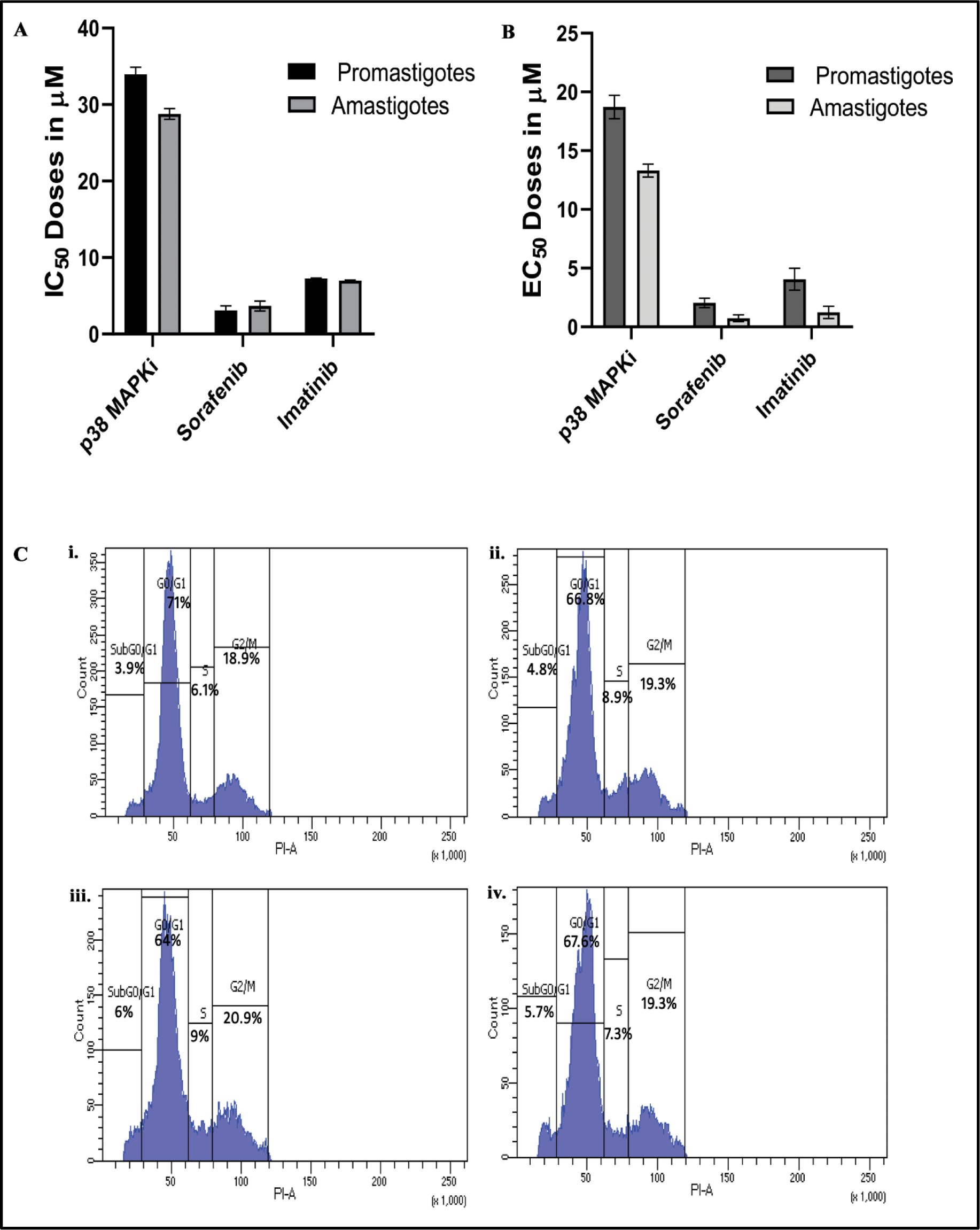
Cell viability assay for 3 inhibitors by MTT. IC_50_ (A) and EC_50_ (B) values are represented by the bar graphs. Data are representatives of atleast three independent experiments. Impact of MAPK inhibitors on cell cycle progression of *L. donovani* promastigotes (C). G1-synchronized log phase promastigote populations, either untreated (i) or treated with IC_50_ doses of p38 MAPK inhibitor IV (ii), Sorafenib (iii) and Imatinib (iv) were analyzed for progression of cell cycle by propidium iodide based FACS analysis (C).

Owing to the information inferred form the above data, role of MAPK in regulating parasite morphology and flagellar length was deciphered by fixing the parasites treated with MAPK inhibitors and visualizing the effects of those inhibitors on parasite morphology by microscopy (Fig. 6). Microscopic images showed that MAPK inhibitors (sorafenib and imatinib, along with the conventional MAPK inhibitor p38 MAPK inhibitor IV) treated cells first transformed into elongated forms in lower doses (1μM) and then lost their promastigote-like shapes and integrity at the IC_50_ doses. No such changes were observed in untreated cells (Fig. 6A-G).

**Figure 6:**
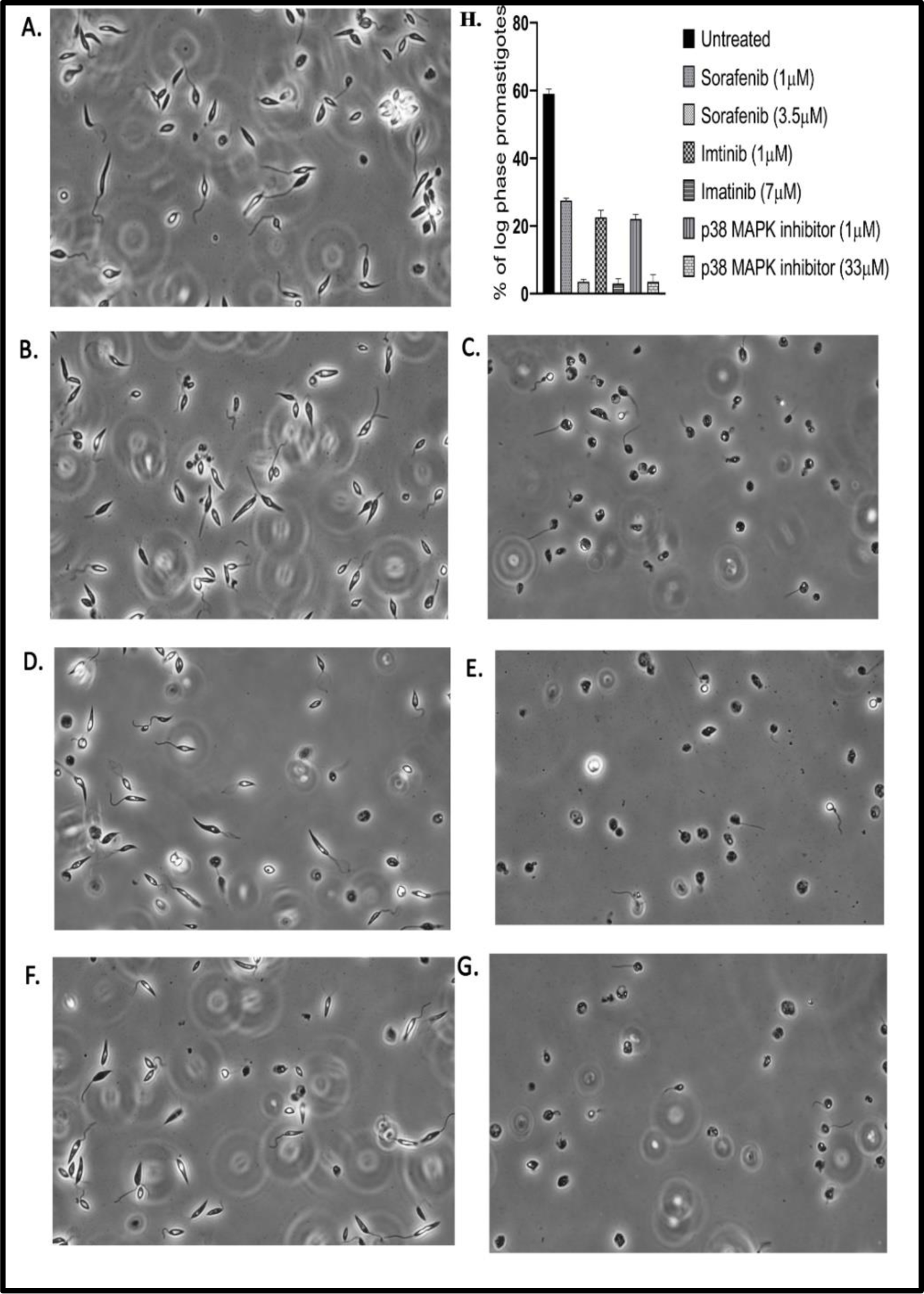
Morphology of *Leishmania donovani* (Ld) promastigotes using phase-contrast microscope. (A) Untreated Ld promastigotes; (B)-(C) Cells treated with Sorafenib (1 μM and 3.5 μM); (D)-(E) Imatinib (1 μM and 7 μM) and (F)-(G) p38 MAPKI IV (1 μM and 33μM). Percentage of log phase promastigotes upon treatment with various inhibitors (H), a schematic representation.

### 3.5. Kinase inhibitors results in the generation of ROS in parasite

ROS generation was studied in order to identify the likely mechanism by which the compound triggered cell death in *Leishmania* promastigotes. The fluorescent microscopy images of H_2_DCFDA ROS are represented after the *Leishmania* promastigotes were exposed to the kinase inhibitors at their IC_50_ doses for 24h, the ROS generation was measured (Fig. 7A). There was a significant increase in ROS generation in p38 MAPK inhibitor, imatinib and sorafenib treated cells (approximately 2-fold increase), compared to the control cells (Fig. 7B).

**Figure 7:**
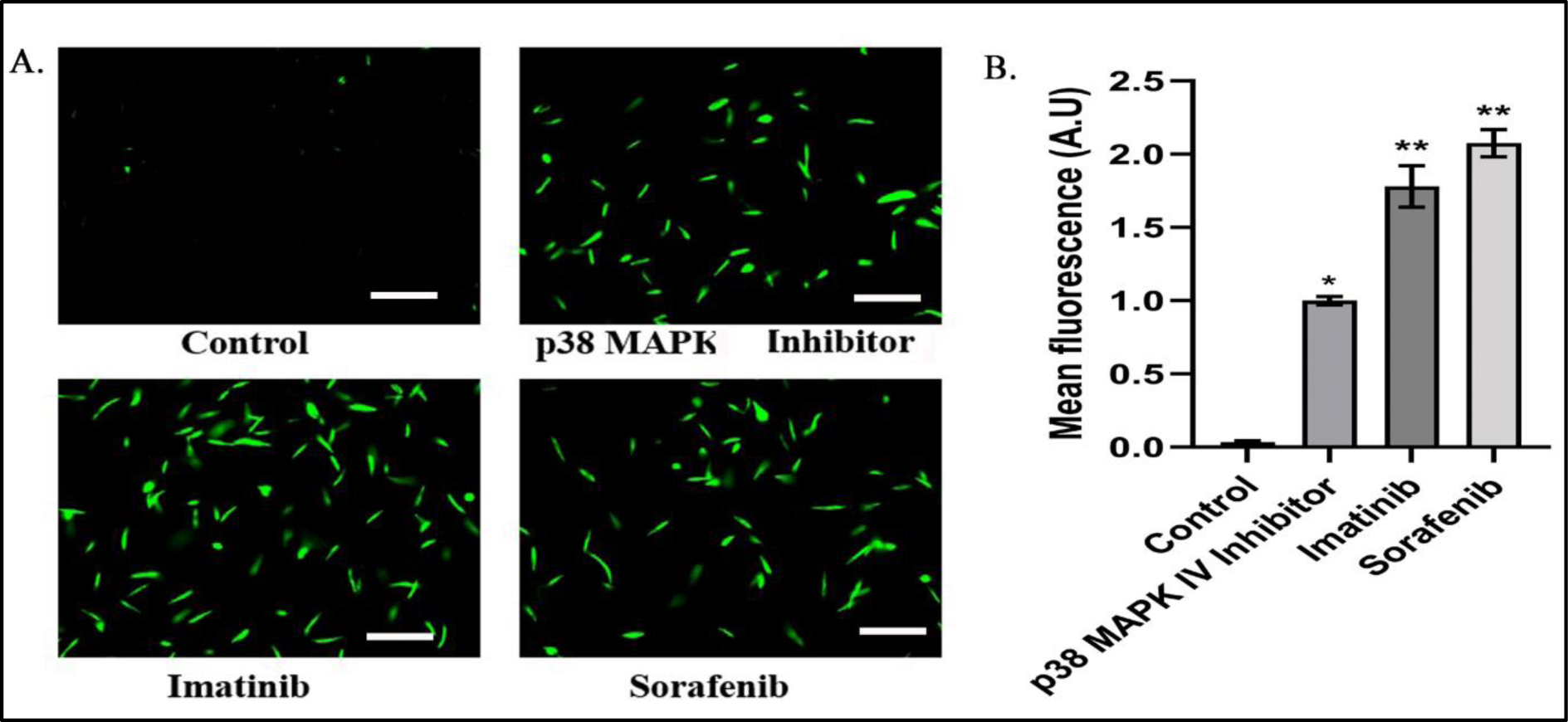
Determination of ROS generation in Ld promastigotes. Ld promastigotes were treated with IC_50_ doses of MAPK inhibitors for 24 hoursand stained with H2DCFDA. ROS generation was observed under fluorescence microscope (scale bar 10 μM) (A). Representative fluorescence intensities of H_2_DCFDA were plotted in the bar diagram after treatment p38 MAPK IV Inhibitor, Imatinib and Sorafenib (B). Results are representative of three individual experiments, and the data represent mean ± SD. *P < 0.05, **P < 0.01 unpaired two tailed t test

### 3.6. Sorafenib and imatinib reduce intra-macrophage parasitic burden and alleviates hepatosplenic parasite load in *Leishmania*-infected Balb/c mice

Since both sorafenib and imatinib showed effects on cell cycle progression and ROS generation, the inhibitors were further explored as anti-leishmanials by analyzing their impacts on parasite multiplication within RAW 264.7 macrophage cell line. The conventional MAPK inhibitor, p38 MAPK inhibitor was used as a control drug. Protein kinase inhibitor treatment resulted in significant decrease in the number of parasites within the macrophages with a maximum effect of 82.0±5.2% reduction at the IC_50_ dose of sorafenib (Fig. 8C). Imatinib also resulted in 62.05±5.2% decrease in the intra-macrophage parasitic burden (Fig. 8D). These results highlight the kinase inhibitors as potential anti-leishmanials.

**Figure 8:**
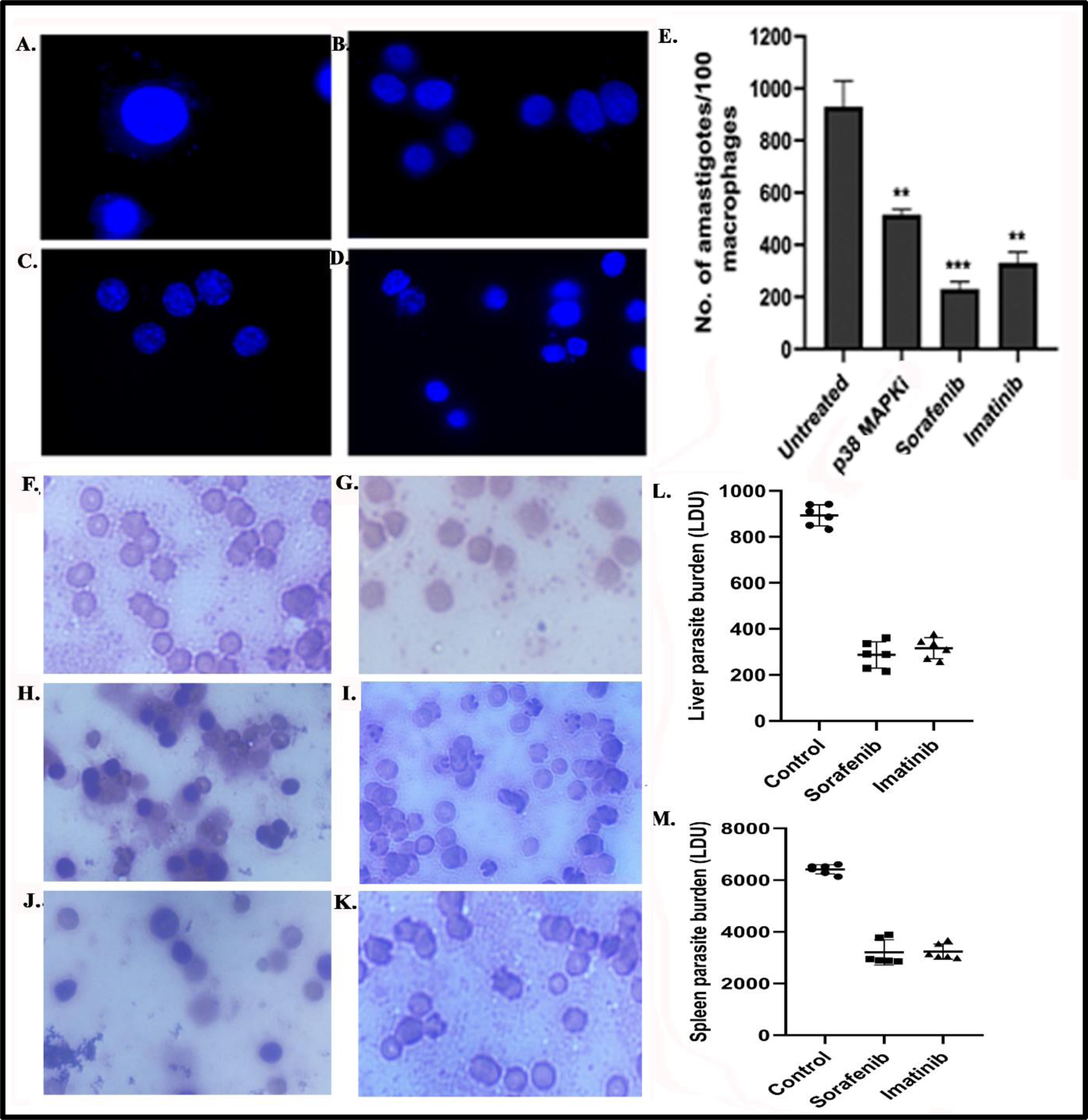
Effects of MAPK inhibitors on parasite survival. Macrophages were infected with Ld log phase promastigotes for 48 hours and then incubated with IC_50_ doses of p38 MAP inhibitor, Sorafenib and Imatinib for another 24 hours. The number of parasites/macrophage was scored for untreated (A), p38 MAP inhibitor (B), Sorafenib (C) and Imatinib (D) by DAPI staining. Representative microscopic image and graphical representation of the same (E) is shown. BALB/c mice were infected with 10^7^ *L. donovani* promastigotes and treated sorafenib and imatinib (10 and 20 mg/kg/day, respectively). The parasite burden in the liver and spleen was determined 45 days after infection and expressed as Leishman–Donovan Units (LDU). Impressions of spleen (control (F), sorafenib-treated (H) and imatinib-treated (J)) and liver (control (G), sorafenib-treated (I) and imatinib-treated (K)) tissues were observed under microscope and disease progression was determined in both and parasite burden expressed as LDU (L and M). Results are representative of three individual experiments, and the data represent mean ± SD. **P < 0.01, *P < 0.05 unpaired two tailed t test.

Subsequently this study focuses on repurposing existing kinase inhibitors as potent anti-leishmanials and owing to the fact that sorafenib and imatinib exhibited anti-proliferative effects on both *Leishmania* promastigotes and amastigotes, they were tested in Balb/c mice to analyse their efficacy in clearing hepatosplenic parasitic burden in *Leishmania*-infected groups. A dose of 10 and 20 mg/kg of body weight of sorafenib and imatinib, respectively, was administered orally for 10 days after 1 month of *Leishmania* infection in Balb/c mice. After the removal of liver and spleen 45 days post infection, multiple impressions of transverse sections of both the organs were prepared for Giemsa staining. Hepato- and splenic parasitic burdens were calculated and expressed as Leishman-Donovan units (LDU) (Fig. 8 L-M). A dose-dependent inhibition was observed at a dose of 10 mg/kg/day for sorafenib with significant reduction of liver and spleen parasitic burden by 74.2 ± 6.2% and 57.79 ± 5.9%, respectively, as compared to untreated *Leishmania*-infected mice. Moreover, similar reduction of parasite burden was also noted when imatinib was administered at a dose of 20 mg/kg/day and liver and spleen parasite burden was reduced by 69.14 ± 5.2% and 46.64 ± 7.1%, respectively (Fig. 8 F-K). Taking these results into consideration, sorafenib and imatinib can be highlighted as anti-leishmanials.

## Conclusion

Protein kinases has evolved into one of the most significant pharmacological targets in the twenty-first century owing to the disruption in the activity of protein kinase in a vast spectrum of diseases, including cancer. The search for orally effective protein kinase inhibitors was spurred by the therapeutic efficacy of imatinib in the treatment of chronic myelogenous leukaemia followed by FDA approval of the drug [34]. Since the approval of imatinib, there has been a significant advancement in the development and discovery of small molecule protein kinase inhibitors [42]. Sixty-nine of the FDA-approved kinase inhibitors have been utilised to treat various types of malignancies, while six others are used to combat inflammatory conditions [22]. Till date, FDA had approved 80 small molecules that are therapeutic protein kinase antagonists, and except three, all of which are efficacious when taken orally [22]. Among such molecules at least five have been enlisted as multi-kinase inhibitors [42]. With increasing reports of development of resistance against tyrosine kinase inhibitors [43], multi-kinase inhibitors are emerging as one of the most preferred regimens [44]. Barring cancer therapeutics, aiming multiple kinase inhibitors might offer an effective strategy against parasitic infections like leishmaniasis. Such approach is crucial in the context of surging reports of resistance against anti-leishmanials including miltefosine, the solitary oral drug available against the infection [45]. With at least 43 essential protein kinases linked to proliferation and infectivity [12], and kinases linked to drug resistance [46] impeding such kinase activity offers a prospective strategy for drug development.

With the paucity of development of selective anti-leishmanials, drug repurposing has long been implemented to treat VL and CL [47]. Classical anti-fungal drugs amphotericin B and pentamidine were introduced as anti-leishmanials [48] and antifungal azoles like fluconazole, econazole have been tested for possible anti-leishmanial potentials [49]. Paramomycin, an aminoglycoside antibiotic primarily applied as drug for ameobiasis also showed promising results as anti-leishmanials [50]. Deleterious side-effects including toxicity and drug resistance remains major impediments in clinical efficacy of the drugs [51] and in this back drop inhibitors of eukaryotic protein kinases (ePK) stem as valid candidates for repurposing.

Though the *Leishmania*-kinome lacks tyrosine kinase and tyrosine kinase like kinases, it comprises at least 195 ePKs belonging to CMGC, AGC, CAMK, STE, NEK, CK1 catalytic domain containing kinases along with *other* kinases including AMPK, Polo, and Aurora kinases [18], [12]. With 23 orthologues, MAP-kinases are the most extensively studied kinases in *Leishmania.* Three of the MAPK orthologues (MKK1, MAPK3, and MAPK9) have been associated with regulation of flagellar length in promastigotes [15]. At least two MAP kinases have been evidenced to modulate the growth, differentiation, and survival of amastigotes [25], [52]. Amino acid and drug transport by aquaglyceroporins (AQP1) are regulated by phosphorylation by a MAPK orthologue MAPK2 [53]. Considering their biological significance in both promastigote and amastigote stages of the parasite, an intense virtual screening was performed with the 23 MAPK orthologues against a library of 12 FDA approved kinase inhibitors. Even though the screening was conducted against to ser-thr-kinases, a hierarchical clustering based on binding affinity identified tyr-kinase inhibitors sorafenib and imatinib, alongside a conventional MAPK inhibitor, p38 MAPK inhibitor IV as potentially antagonistic for multiple MAPKs with binding energy ranging between -6.4 kcal/mol to -9.2 kcal/mol. Sorafenib is a multikinase inhibitor with the potential to aim both ser-thr kinases and receptor tyr-kinases [39]. Imatinib selectively targets non-receptor tyrosine kinases like BCR-Abl and c-Src [54] and was evidenced to be effective in *L. major* CL model in BALB/c mice and *L. amazonensis* infected mice earlier [55]. Both the inhibitors displayed highest affinity for LdMAP-kinase (LdBPK_323450.1_ *Leishmania donovani* BPK282A1) with binding energies of -8.9 Kcal/mol and -10.2 Kcal/mol, respectively. Albeit the binding modes differs compared to the typical binding pockets in the mammalian targets for the inhibitors, divulging stability of the complexes by MD simulation projected stable complex formation with in the kinase conserved domains. A comparative ADMET analysis revealed that both sorafenib and imatinib satisfy Lipinski drug-likeness, and greater GI absorption rate compared to the approved drugs, amphotericin B and miltefosine highlighting both sorafenib and imatinib as potent repurposed drug candidates against leishmaniasis. With an EC_50_ value of 3.5 μM against promastigotes and 3.12 μM against amastigotes, sorafenib demonstrated profound antileishmanial action. The observation also corroborated *in vivo* with experimental models of VL in BALB/c mice with significant abrogation of parasitic load in spleen and liver. In recent years drug repurposing against *Leishmania*, or against trypanosomatids *per se*, offered silver lines in prevailing over drug non-responsiveness. Combining miltefosine with the antitumoral selective estrogen receptor modulator tamoxifen can overcome miltefosine resistance in in vivo experimental models of CL [56]. Deeper preclinical studies with sorafenib and imatinib as single or combinatorial therapeutics with available drugs would enrich the existing repertoire of anti=leishmanials.

## Data availability statement

All datasets generated for this study are included in the article/Supplementary Material.

## Ethics statement

The animal study was approved by the Institutional Animal Ethics Committee on Animal Experiments of the University of Kalyani (892/GO/Re/S/01/CPCSEA). The maintenance and culture of *L. donovani* was also in strict regulations according to the recommendations in the Institutional Biosafety Committee (Ref No. IBSC-KU/2023/A4), Kalyani University (IBSC-KU).

## Author contributions

Conceptualization: AruB and AriB. Data curation: AniB, ArB and SS. Formal analysis: AruB, AriB and AniB. Funding acquisition: AruB and AniB. Investigation: AniB, SS, SB, ArB. Methodology: AriB, AruB and AniB. Project administration: AruB and AriB. Resources: AruB and AriB. Software: AniB, ArB and AriB. Supervision: AruB and AriB. Validation: AniB, AruB and AriB. Visualization: AniB. A Roles/Writing - original draft: AniB and Writing - review & editing: AruB and AriB.

## Acknowledgements

The authors would like to acknowledge the DST-PURSE Programme and PRG, University of Kalyani, for the funding. Authors acknowledge the contribution of Mr. Tamal Ghosh, IISER, Kolkata for performing the flow cytometry experiments, Dr. Kumaresh Ghosh, Professor, Department of Chemistry, University of Kalyani, for allowing to use the fluorescence microscope of his laboratory and Dr. Analabha Roy, Assistant Professor, Department of Physics, The University of Burdwan for his contribution in generating MD simulation data.

## References

[1] A. Biswas, A. Bhattacharjee, P. K. Das, A. Biswas, A. Bhattacharjee, and P. K. Das, “Role of cAMP Homeostasis in Intra-Macrophage Survival and Infectivity of Unicellular Parasites like *Leishmania*,” in Vector-Borne Diseases - Recent Developments in Epidemiology and Control, IntechOpen, 2019. doi: 10.5772/intechopen.86360.

[2] D. Rosenzweig, D. Smith, P. J. Myler, R. W. Olafson, and D. Zilberstein, “Post-translational modification of cellular proteins during Leishmania donovani differentiation,” Proteomics, vol. 8, no. 9, pp. 1843–1850, May 2008, doi: 10.1002/pmic.200701043.

[3] J. Joshi, N. Malla, and S. Kaur, “A comparative evaluation of efficacy of chemotherapy, immunotherapy and immunochemotherapy in visceral leishmaniasis-an experimental study,” Parasitology International, vol. 63, no. 4, pp. 612–620, Aug. 2014, doi: 10.1016/j.parint.2014.04.002.

[4] J. Tanwar, S. Das, Z. Fatima, and S. Hameed, “Multidrug Resistance: An Emerging Crisis,” Interdisciplinary Perspectives on Infectious Diseases, vol. 2014, p. e541340, Jul. 2014, doi: 10.1155/2014/541340.

[5] R. J. Wheeler, E. Gluenz, and K. Gull, “The cell cycle of Leishmania: morphogenetic events and their implications for parasite biology,” Mol Microbiol, vol. 79, no. 3, pp. 647–662, Feb. 2011, doi: 10.1111/j.1365-2958.2010.07479.x.

[6] T. Garcia-Garcia, S. Poncet, A. Derouiche, L. Shi, I. Mijakovic, and M.-F. Noirot-Gros, “Role of Protein Phosphorylation in the Regulation of Cell Cycle and DNA-Related Processes in Bacteria,” Frontiers in Microbiology, vol. 7, 2016, Accessed: Jan. 06, 2024. [Online]. Available: https://www.frontiersin.org/articles/10.3389/fmicb.2016.00184

[7] A. Efstathiou and D. Smirlis, “Leishmania Protein Kinases: Important Regulators of the Parasite Life Cycle and Molecular Targets for Treating Leishmaniasis,” Microorganisms, vol. 9, no. 4, p. 691, Mar. 2021, doi: 10.3390/microorganisms9040691.

[8] M. Parsons, M. Valentine, and V. Carter, “Protein kinases in divergent eukaryotes: identification of protein kinase activities regulated during trypanosome development,” Proc. Natl. Acad. Sci. U.S.A., vol. 90, no. 7, pp. 2656–2660, Apr. 1993, doi: 10.1073/pnas.90.7.2656.

[9] M. Parsons, M. Valentine, J. Deans, G. L. Schieven, and J. A. Ledbetter, “Distinct patterns of tyrosine phosphorylation during the life cycle of Trypanosoma brucei,” Mol. Biochem. Parasitol., vol. 45, no. 2, pp. 241–248, Apr. 1991, doi: 10.1016/0166-6851(91)90091-j.

[10] E. Wheeler-Alm and S. Z. Shapiro, “Evidence of tyrosine kinase activity in the protozoan parasite Trypanosoma brucei,” J. Protozool., vol. 39, no. 3, pp. 413–416, Jun. 1992, doi: 10.1111/j.1550-7408.1992.tb01473.x.

[11] E. S. Nakayasu, M. R. Gaynor, T. J. P. Sobreira, J. A. Ross, and I. C. Almeida, “Phosphoproteomic analysis of the human pathogen *Trypanosoma cruzi* at the epimastigote stage,” Proteomics, vol. 9, no. 13, pp. 3489–3506, Jul. 2009, doi: 10.1002/pmic.200800874.

[12] B. N, et al., “Systematic functional analysis of Leishmania protein kinases identifies regulators of differentiation or survival,” bioRxiv, p. 2020.09.06.279091, Dec. 2020, doi: 10.1101/2020.09.06.279091.

[13] M. Wiese, “Leishmania MAP kinases – Familiar proteins in an unusual context,” International Journal for Parasitology, vol. 37, no. 10, pp. 1053–1062, Aug. 2007, doi: 10.1016/j.ijpara.2007.04.008.

[14] M. Wiese, D. Kuhn, and C. G. Grünfelder, “Protein kinase involved in flagellar-length control,” Eukaryot Cell, vol. 2, no. 4, pp. 769–777, Aug. 2003, doi: 10.1128/EC.2.4.769-777.2003.

[15] F. Bengs, A. Scholz, D. Kuhn, and M. Wiese, “LmxMPK9, a mitogen-activated protein kinase homologue affects flagellar length in Leishmania mexicana,” Mol Microbiol, vol. 55, no. 5, pp. 1606–1615, Mar. 2005, doi: 10.1111/j.1365-2958.2005.04498.x.

[16] D. Kuhn and M. Wiese, “LmxPK4, a mitogen-activated protein kinase kinase homologue of Leishmania mexicana with a potential role in parasite differentiation,” Mol Microbiol, vol. 56, no. 5, pp. 1169–1182, Jun. 2005, doi: 10.1111/j.1365-2958.2005.04614.x.

[17] M. Erdmann, A. Scholz, I. M. Melzer, C. Schmetz, and M. Wiese, “Interacting protein kinases involved in the regulation of flagellar length,” Mol Biol Cell, vol. 17, no. 4, pp. 2035–2045, Apr. 2006, doi: 10.1091/mbc.e05-10-0976.

[18] M. Parsons, E. A. Worthey, P. N. Ward, and J. C. Mottram, “Comparative analysis of the kinomes of three pathogenic trypanosomatids: Leishmania major, Trypanosoma brucei and Trypanosoma cruzi,” BMC Genomics, vol. 6, p. 127, Sep. 2005, doi: 10.1186/1471-2164-6-127.

[19] M. Wiese, “A mitogen-activated protein (MAP) kinase homologue of Leishmania mexicana is essential for parasite survival in the infected host,” EMBO J, vol. 17, no. 9, pp. 2619–2628, May 1998, doi: 10.1093/emboj/17.9.2619.

[20] P. Kaur, M. Garg, A. Hombach-Barrigah, J. Clos, and N. Goyal, “MAPK1 of Leishmania donovani interacts and phosphorylates HSP70 and HSP90 subunits of foldosome complex,” Sci Rep, vol. 7, no. 1, p. 10202, Aug. 2017, doi: 10.1038/s41598-017-09725-w.

[21] M. A. Morales, O. Renaud, W. Faigle, S. L. Shorte, and G. F. Späth, “Over-expression of Leishmania major MAP kinases reveals stage-specific induction of phosphotransferase activity,” Int J Parasitol, vol. 37, no. 11, pp. 1187–1199, Sep. 2007, doi: 10.1016/j.ijpara.2007.03.006.

[22] R. Roskoski, “Properties of FDA-approved small molecule protein kinase inhibitors: A 2024 update,” Pharmacol Res, vol. 200, p. 107059, Jan. 2024, doi: 10.1016/j.phrs.2024.107059.

[23] K. H. Kim and J. M. Sederstrom, “Assaying cell cycle status using flow cytometry,” Curr Protoc Mol Biol, vol. 111, p. 28.6.1–28.6.11, Jul. 2015, doi: 10.1002/0471142727.mb2806s111.

[24] N. M. O’Boyle, M. Banck, C. A. James, C. Morley, T. Vandermeersch, and G. R. Hutchison, “Open Babel: An open chemical toolbox,” J Cheminform, vol. 3, p. 33, Oct. 2011, doi: 10.1186/1758-2946-3-33.

[25] M. Cayla et al., “Transgenic Analysis of the Leishmania MAP Kinase MPK10 Reveals an Auto-inhibitory Mechanism Crucial for Stage-Regulated Activity and Parasite Viability,” PLoS Pathog, vol. 10, no. 9, p. e1004347, Sep. 2014, doi: 10.1371/journal.ppat.1004347.

[26] P. Cohen, “Targeting protein kinases for the development of anti-inflammatory drugs,” Curr Opin Cell Biol, vol. 21, no. 2, pp. 317–324, Apr. 2009, doi: 10.1016/j.ceb.2009.01.015.

[27] B. B. Friday and A. A. Adjei, “Advances in targeting the Ras/Raf/MEK/Erk mitogen-activated protein kinase cascade with MEK inhibitors for cancer therapy,” Clin Cancer Res, vol. 14, no. 2, pp. 342–346, Jan. 2008, doi: 10.1158/1078-0432.CCR-07-4790.

[28] T. R. Rheault et al., “Discovery of Dabrafenib: A Selective Inhibitor of Raf Kinases with Antitumor Activity against B-Raf-Driven Tumors,” ACS Med Chem Lett, vol. 4, no. 3, pp. 358– 362, Feb. 2013, doi: 10.1021/ml4000063.

[29] Y. Lee, K. J. Bae, H. J. Chon, S. H. Kim, S. A. Kim, and J. Kim, “A Receptor Tyrosine Kinase Inhibitor, Dovitinib (TKI-258), Enhances BMP-2-Induced Osteoblast Differentiation In Vitro,” Mol Cells, vol. 39, no. 5, pp. 389–394, May 2016, doi: 10.14348/molcells.2016.2300.

[30] Z. T. Al-Salama and S. J. Keam, “Entrectinib: First Global Approval,” Drugs, vol. 79, no. 13, pp. 1477–1483, Sep. 2019, doi: 10.1007/s40265-019-01177-y.

[31] M. A. S. Abourehab, A. M. Alqahtani, B. G. M. Youssif, and A. M. Gouda, “Globally Approved EGFR Inhibitors: Insights into Their Syntheses, Target Kinases, Biological Activities, Receptor Interactions, and Metabolism,” Molecules, vol. 26, no. 21, p. 6677, Nov. 2021, doi: 10.3390/molecules26216677.

[32] J. Sharifi-Rad et al., “Genistein: An Integrative Overview of Its Mode of Action, Pharmacological Properties, and Health Benefits,” Oxid Med Cell Longev, vol. 2021, p. 3268136, Jul. 2021, doi: 10.1155/2021/3268136.

[33] I. E. Ahn and J. R. Brown, “Targeting Bruton’s Tyrosine Kinase in CLL,” Front. Immunol., vol. 12, p. 687458, Jun. 2021, doi: 10.3389/fimmu.2021.687458.

[34] P. Cohen, D. Cross, and P. A. Jänne, “Kinase drug discovery 20 years after imatinib: progress and future directions,” Nat Rev Drug Discov, vol. 20, no. 7, Art. no. 7, Jul. 2021, doi: 10.1038/s41573-021-00195-4.

[35] S. R. D. Johnston and A. Leary, “Lapatinib: a novel EGFR/HER2 tyrosine kinase inhibitor for cancer,” Drugs Today (Barc), vol. 42, no. 7, pp. 441–453, Jul. 2006, doi: 10.1358/2006.42.7.985637.

[36] B. R. Davies et al., “Preclinical Pharmacology of AZD5363, an Inhibitor of AKT: Pharmacodynamics, Antitumor Activity, and Correlation of Monotherapy Activity with Genetic Background,” Molecular Cancer Therapeutics, vol. 11, no. 4, pp. 873–887, Apr. 2012, doi: 10.1158/1535-7163.MCT-11-0824-T.

[37] P. Vuylsteke et al., “Pictilisib PI3Kinase inhibitor (a phosphatidylinositol 3-kinase [PI3K] inhibitor) plus paclitaxel for the treatment of hormone receptor-positive, HER2-negative, locally recurrent, or metastatic breast cancer: interim analysis of the multicentre, placebo-controlled, phase II randomised PEGGY study,” Ann Oncol, vol. 27, no. 11, pp. 2059–2066, Nov. 2016, doi: 10.1093/annonc/mdw320.

[38] F. H. Tan, T. L. Putoczki, S. S. Stylli, and R. B. Luwor, “Ponatinib: a novel multi-tyrosine kinase inhibitor against human malignancies,” Onco Targets Ther, vol. 12, pp. 635–645, Jan. 2019, doi: 10.2147/OTT.S189391.

[39] S. M. Wilhelm, L. Adnane, P. Newell, A. Villanueva, J. M. Llovet, and M. Lynch, “Preclinical overview of sorafenib, a multikinase inhibitor that targets both Raf and VEGF and PDGF receptor tyrosine kinase signaling,” Mol Cancer Ther, vol. 7, no. 10, pp. 3129–3140, Oct. 2008, doi: 10.1158/1535-7163.MCT-08-0013.

[40] E. Liu, N. Aslam, G. Nigam, and J. K. Limdi, “Tofacitinib and newer JAK inhibitors in inflammatory bowel disease—where we are and where we are going,” Drugs Context, vol. 11, pp. 2021–11–4, Apr. 2022, doi: 10.7573/dic.2021-11-4.

[41] A. Daina, O. Michielin, and V. Zoete, “SwissADME: a free web tool to evaluate pharmacokinetics, drug-likeness and medicinal chemistry friendliness of small molecules,” Sci Rep, vol. 7, p. 42717, Mar. 2017, doi: 10.1038/srep42717.

[42] K. S. Bhullar et al., “Kinase-targeted cancer therapies: progress, challenges and future directions,” Mol Cancer, vol. 17, p. 48, Feb. 2018, doi: 10.1186/s12943-018-0804-2.

[43] Y. Yang, S. Li, Y. Wang, Y. Zhao, and Q. Li, “Protein tyrosine kinase inhibitor resistance in malignant tumors: molecular mechanisms and future perspective,” Signal Transduct Target Ther, vol. 7, no. 1, p. 329, Sep. 2022, doi: 10.1038/s41392-022-01168-8.

[44] Y.-L. Lai, K.-H. Wang, H.-P. Hsieh, and W.-C. Yen, “Novel FLT3/AURK multikinase inhibitor is efficacious against sorafenib-refractory and sorafenib-resistant hepatocellular carcinoma,” J Biomed Sci, vol. 29, no. 1, p. 5, Jan. 2022, doi: 10.1186/s12929-022-00788-0.

[45] A. Bhattacharya and M. Ouellette, “New insights with miltefosine unresponsiveness in Brazilian Leishmania infantum isolates,” EBioMedicine, vol. 37, pp. 13–14, Nov. 2018, doi: 10.1016/j.ebiom.2018.10.016.

[46] A. Bhattacharya et al., “Coupling chemical mutagenesis to next generation sequencing for the identification of drug resistance mutations in Leishmania,” Nat Commun, vol. 10, no. 1, p. 5627, Dec. 2019, doi: 10.1038/s41467-019-13344-6.

[47] M. Berriman et al., “The Genome of the African Trypanosome Trypanosoma brucei,” Science, vol. 309, no. 5733, pp. 416–422, Jul. 2005, doi: 10.1126/science.1112642.

[48] S. Sundar and J. Chakravarty, “Liposomal amphotericin B and leishmaniasis: dose and response,” J Glob Infect Dis, vol. 2, no. 2, pp. 159–166, May 2010, doi: 10.4103/0974-777X.62886.

[49] S. Emami, P. Tavangar, and M. Keighobadi, “An overview of azoles targeting sterol 14α-demethylase for antileishmanial therapy,” Eur J Med Chem, vol. 135, pp. 241–259, Jul. 2017, doi: 10.1016/j.ejmech.2017.04.044.

[50] N. Sosa et al., “Topical paromomycin for New World cutaneous leishmaniasis,” PLoS Negl Trop Dis, vol. 13, no. 5, p. e0007253, May 2019, doi: 10.1371/journal.pntd.0007253.

[51] F. Frézard et al., “Liposomal Amphotericin B for Treatment of Leishmaniasis: From the Identification of Critical Physicochemical Attributes to the Design of Effective Topical and Oral Formulations,” Pharmaceutics, vol. 15, no. 1, p. 99, Dec. 2022, doi: 10.3390/pharmaceutics15010099.

[52] S. M. Duncan et al., “Conditional gene deletion with DiCre demonstrates an essential role for CRK3 in Leishmania mexicana cell cycle regulation,” Mol Microbiol, vol. 100, no. 6, pp. 931–944, Jun. 2016, doi: 10.1111/mmi.13375.

[53] G. Mandal et al., “Modulation of Leishmania major aquaglyceroporin activity by a mitogen-activated protein kinase,” Mol Microbiol, vol. 85, no. 6, pp. 1204–1218, Sep. 2012, doi: 10.1111/j.1365-2958.2012.08169.x.

[54] Y. Tsutsui, D. Deredge, P. L. Wintrode, and F. A. Hays, “Imatinib binding to human c-Src is coupled to inter-domain allostery and suggests a novel kinase inhibition strategy,” Sci Rep, vol. 6, p. 30832, Aug. 2016, doi: 10.1038/srep30832.

[55] M. Moslehi et al., “Study of therapeutic effect of different concentrations of imatinib on Balb/c model of cutaneous leishmaniasis,” AIMS Microbiol, vol. 6, no. 2, pp. 152–161, 2020, doi: 10.3934/microbiol.2020010.

[56] C. T. Trinconi, J. Q. Reimão, A. C. Coelho, and S. R. B. Uliana, “Efficacy of tamoxifen and miltefosine combined therapy for cutaneous leishmaniasis in the murine model of infection with Leishmania amazonensis,” J Antimicrob Chemother, vol. 71, no. 5, pp. 1314–1322, May 2016, doi: 10.1093/jac/dkv495.

