## supplementary for "Repurposing approved protein kinase inhibitors as potent anti-leishmanials targeting *Leishmania* MAP kinases"

Supplementary data

Figure S1: Ld MAP kinase protein models and their corresponding Ramachandran plots

Figure S2: *Leishmania donovani* MAP Kinase, LdBPK-323450.1_ *Leishmania donovani*-BPK282A1 interactions with inhibitors

Figure S3: Molecular docking of *Leishmania donovani* MAP Kinase, LdBPK-323450.1_ *Leishmania donovani*-BPK282A1 with entrectinib and ponatinib.

Table S1: Interaction data for each binding site of 3 inhibitors in complex with *Leishmania donovani* MAP Kinase, LdBPK-323450.1_ *Leishmania donovani*-BPK282A1.

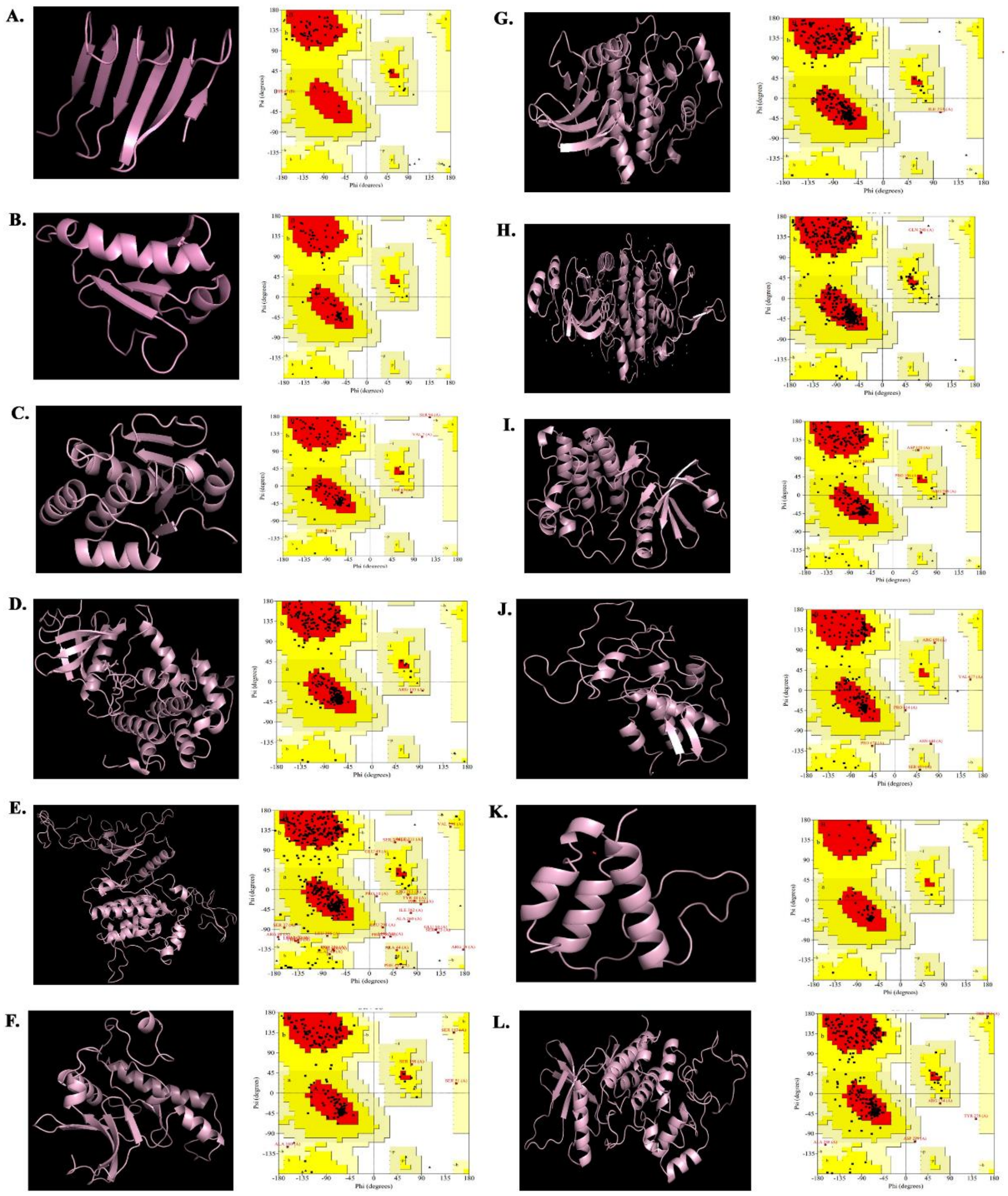

A.
B.
C.
D.
E.
F.
G.
H.
I.
J.
K.
L.
Phi (degrees)
Psi (degrees)

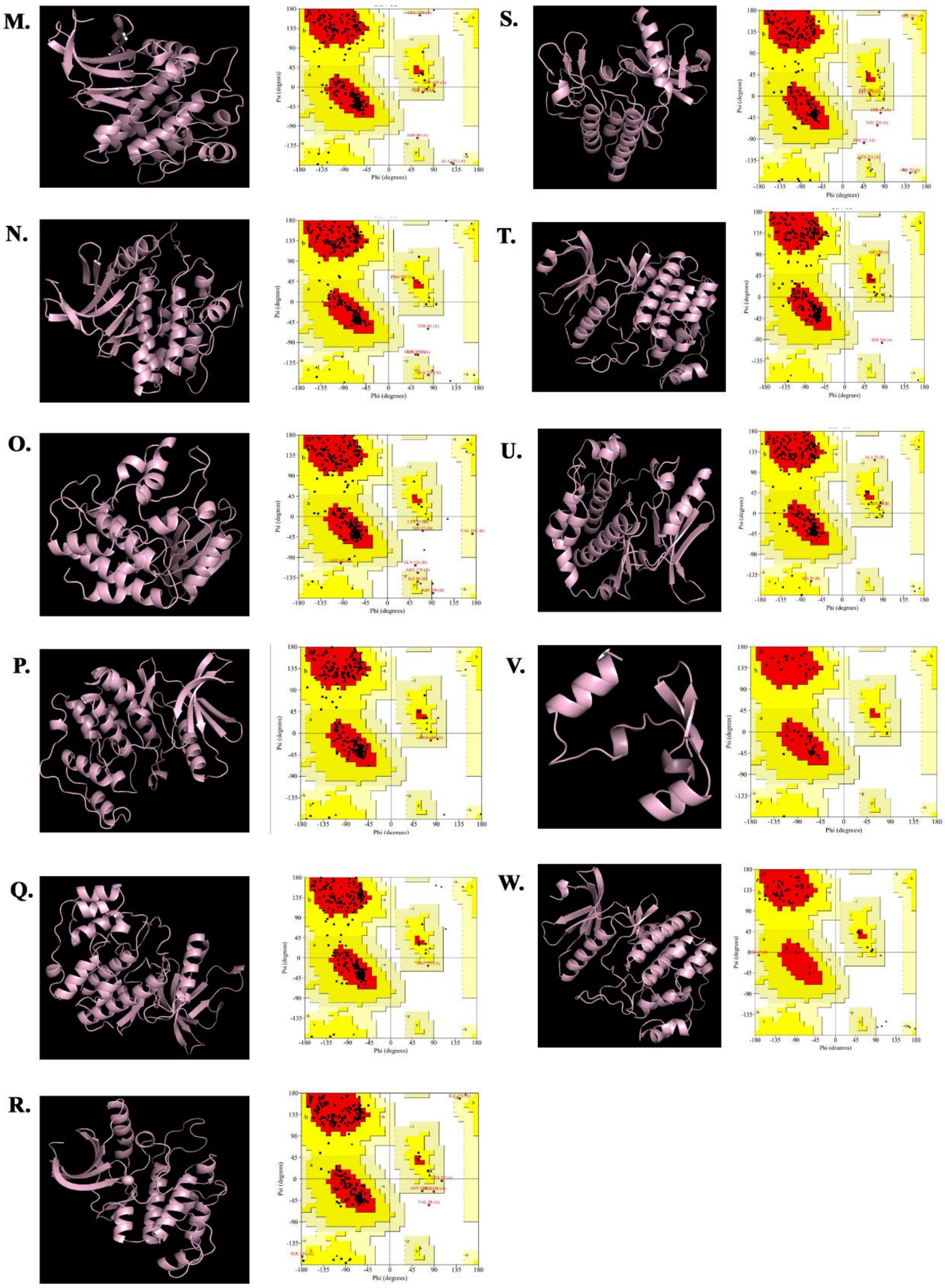


Figure S1: Protein models were built using Swiss-Modelers for all the 23 Ld MAPKs and corresponding Ramachandran plots were derived from Procheck and models with 0-1 amino acids in the questionable zone were selected for further studies.

| IMATINIB | | |
|---|---|---|
| Hydrophobic Interactions | **RESIDUE** | **AA** |
| | 10A | LEU |
| | 10A | LEU |
| | 10A | LEU |
| | 18A | VAL |
| | 82A | PHE |
| Hydrogen Bonds | 14A | THR |
| | 15A | TYR |
| | 16A | GLY |
| | 83A | ILE |
| | 131A | ASN |
| | 131A | ASN |
| **SORAFENIB** | | |
| Hydrophobic Interactions | **RESIDUE** | **AA** |
| | 10A | LEU |
| | 82A | PHE |
| | 133A | LEU |
| **p38 MAPK INHIBITOR IV** | | |
| Hydrophobic Interactions | **RESIDUE** | **AA** |
| | 10A | LEU |
| | 10A | LEU |
| | 10A | LEU |
| | 18A | VAL |
| | 31A | ALA |
| | 64A | ILE |
| | 80A | PHE |
| | 89A | GLN |
| | 133A | LEU |
| | 133A | LEU |
| | 83A | ILE |
| | 83A | ILE |
| | 83A | ASP |

Table S1: Interaction data (Hydrophobic Interactions and Hydrogen Bonds) is provided for each binding site for 3 complexes. (A) LdMAPK-imatinib complex, (B) LdMAPK-sorafenib complex, and (C) LdMAPK-p38 inhibitor IV complex.

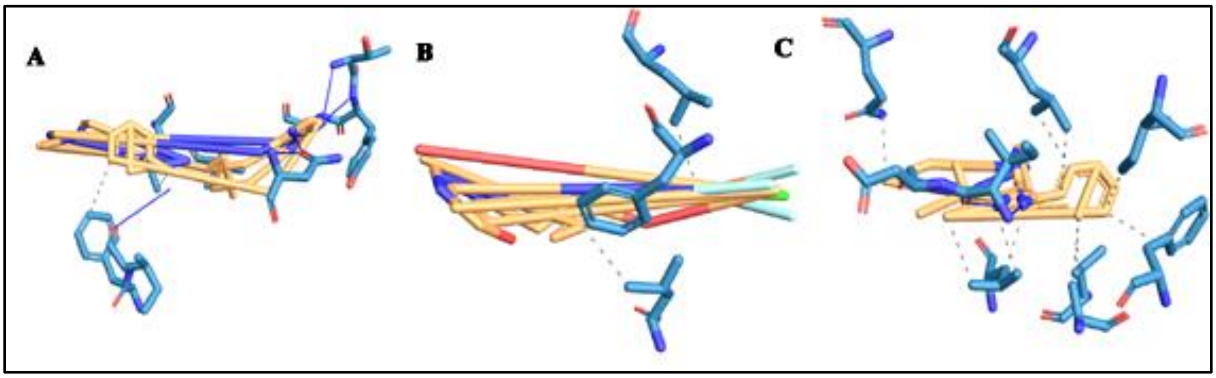

Figure S2: *Leishmania donovani* MAP Kinase, LdBPK-323450.1_ *Leishmania donovani*-BPK282A1 with different inhibitors. (A) Imatinib, (B) Sorafenib, (C) p58 MAP Kinase inhibitor. Large parts of all ligands bind via hydrophobic interactions

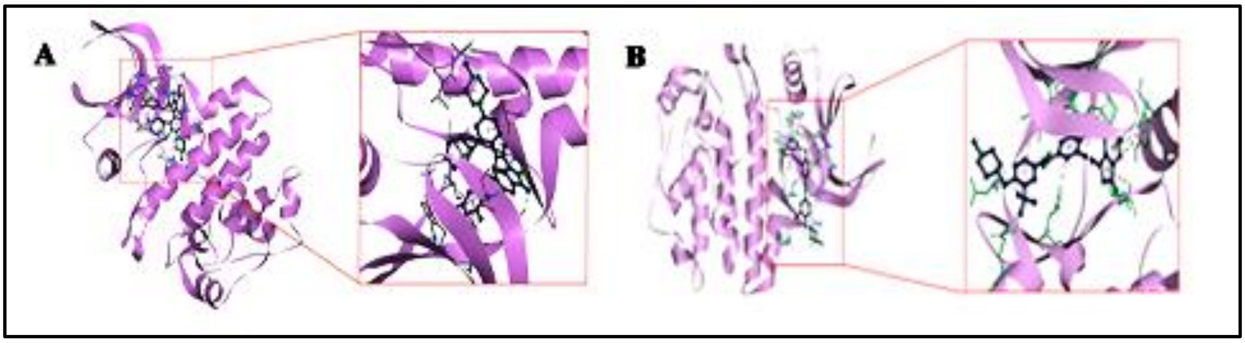

Figure S3: Interactions, bond lengths and hydrogen bond donor and acceptor around ligand binding pocket for Ld MAPK-ligand interaction are shown in these figures. Interaction of LdBPK_323450.1_ *Leishmania donovani* BPK282A1 with entrectinib (A) and ponatinib (B).
